# SpatialQuery: scalable discovery and molecular characterization of multicellular motifs from spatial omics data

**DOI:** 10.64898/2026.04.22.720136

**Authors:** Shaokun An, Mark Keller, Nils Gehlenborg, Martin Hemberg

**Affiliations:** Gene Lay Institute of Immunology and Inflammation, Brigham and Women’s Hospital, Massachusetts General Hospital and Harvard Medical School, Boston, MA, USA; Department of Biomedical Informatics, Harvard Medical School, Boston, MA, USA

## Abstract

Spatially resolved single-cell technologies enable profiling of cells *in situ*, yet computational approaches that jointly discover multicellular spatial patterns and characterize their molecular programs remain limited. Here we introduce SpatialQuery, a framework that can both identify cellular motifs, i.e. recurrent multicellular co-localization patterns, and perform molecular analyses focused on the motifs. It uncovers genes modulated by spatial contexts through differential expression analysis, and detects coordinated expression changes through covariation analysis. SpatialQuery can identify functional tissue units, and goes beyond pairwise analyses to characterize multicellular interactions. Applications to both spatial transcriptomics and proteomics data uncover cross-germ-layer signaling in gut tube patterning, disease-specific fibrotic and immunosuppressive niches in kidney and colon, and regional determinants of motif-associated transcriptional programs in a mouse brain atlas. SpatialQuery is available as a Python package, and we demonstrate how its light computational footprint enables integration into web-based cell atlas portals for interactive visualization and exploration.

## Introduction

Spatially resolved single-cell technologies, including imaging-based spatial transcriptomics (e.g. MERFISH, seqFISH, CosMx, and Xenium) and spatial proteomics platforms (e.g. CODEX, MIBI, and CellDIVE), enable simultaneous profiling of molecular states while retaining spatial context for millions of cells in a single experiment^1,2^. These advances have enabled new directions of research to decipher how cells are organized *in situ* and how their microenvironment affects molecular identity. A central question that emerges is how specific combinations of neighboring cell types shape the transcriptional or proteomic state, and how they impact tissue function. Addressing this question requires two complementary analytical tasks: identifying which cell type combinations recurrently co-localize beyond random expectation, and characterizing the molecular programs associated with these multicellular spatial configurations^3^.

Several computational methods have been developed for characterizing spatial cellular neighborhoods. One approach has been spatial domain identification, which seeks to segment tissues into transcriptionally or compositionally coherent regions^4–8^, while niche-focused methods seek to identify fine grained spatial structures^9–14^. These methods have advanced the ability to define distinct cellular communities, and in several cases it has been possible to associate domains with gene programs or specific cell-cell interactions^9,15^. Nonetheless, the identification of spatial neighborhoods and the molecular characterization of cells within neighborhoods are often treated as separate analytical tasks. They are typically addressed by distinct tools with different assumptions, input requirements, and computational frameworks. The decoupling of tools introduces inconsistent cell groups across analytical stages, preventing the molecular analysis from fully leveraging the identified spatial patterns.

Methods for characterizing molecular relationships between spatially proximal cells have advanced along two largely independent lines. One line of investigation addresses how gene-gene correlations within cells vary across space^16–19^, or seek to remove spatial confounders to recover such correlations^20,21^. These methods reveal how gene regulatory programs within a cell are spatially organized. A second line concerns molecular coordination between neighboring cells, based on the expression of ligand-receptor (LR) pairs in combination with spatial distance constraints^22–28^, with or without the use of a curated database of interactions^29–32^. Despite the methodological advances, several challenges remain. Most methods for inferring cell-cell communication consider only pairwise interactions^33^, and since they implicitly assume that the tissue is homogenous to aggregate signals, the role of the microenvironment is ignored. Moreover, many current approaches depend on computationally intensive procedures constraining real-time exploratory analysis and limiting scalability^10,22,34^.

Here we present SpatialQuery, a computational framework that addresses these challenges by unifying multicellular spatial motif discovery with motif-associated molecular characterization from spatially resolved single-cell data, including spatial transcriptomics and spatial proteomics platforms (Fig. 1). We define a cellular motif as a recurrent, statistically enriched group of co-localized cell types, distinct from domains that segment tissue into spatially coherent regions, or niches that classify neighborhoods by compositional similarity. The framework provides three integrated analytical modules which extend beyond pairwise cell-type analyses: (1) motif enrichment analysis to discover and statistically assess cellular motifs, (2) motif-associated differential expression analysis to identify genes modulated by specific neighborhood contexts, and (3) motif-associated cross-cell covariation analysis to detect gene pairs exhibiting coordinated expression changes within a motif. SpatialQuery achieves computational efficiency through closed-form statistical tests and efficient data structures, enabling atlas-scale analysis of millions of cells within minutes. SpatialQuery supports both single-sample and multi-sample analyses, and for the latter it enables both condition-specific and condition-stratified analyses. We demonstrate the utility of SpatialQuery across four biological systems and spatial platforms: mouse embryo gut tube patterning (seqFISH)^35^, kidney disease microenvironments (Slide-seqV2)^36^, colorectal cancer immunosuppressive niches (CODEX)^37^ and a whole-brain MERFISH atlas comprising ∼8.4 million cells across 239 tissue sections^38^.

**Fig. 1.**
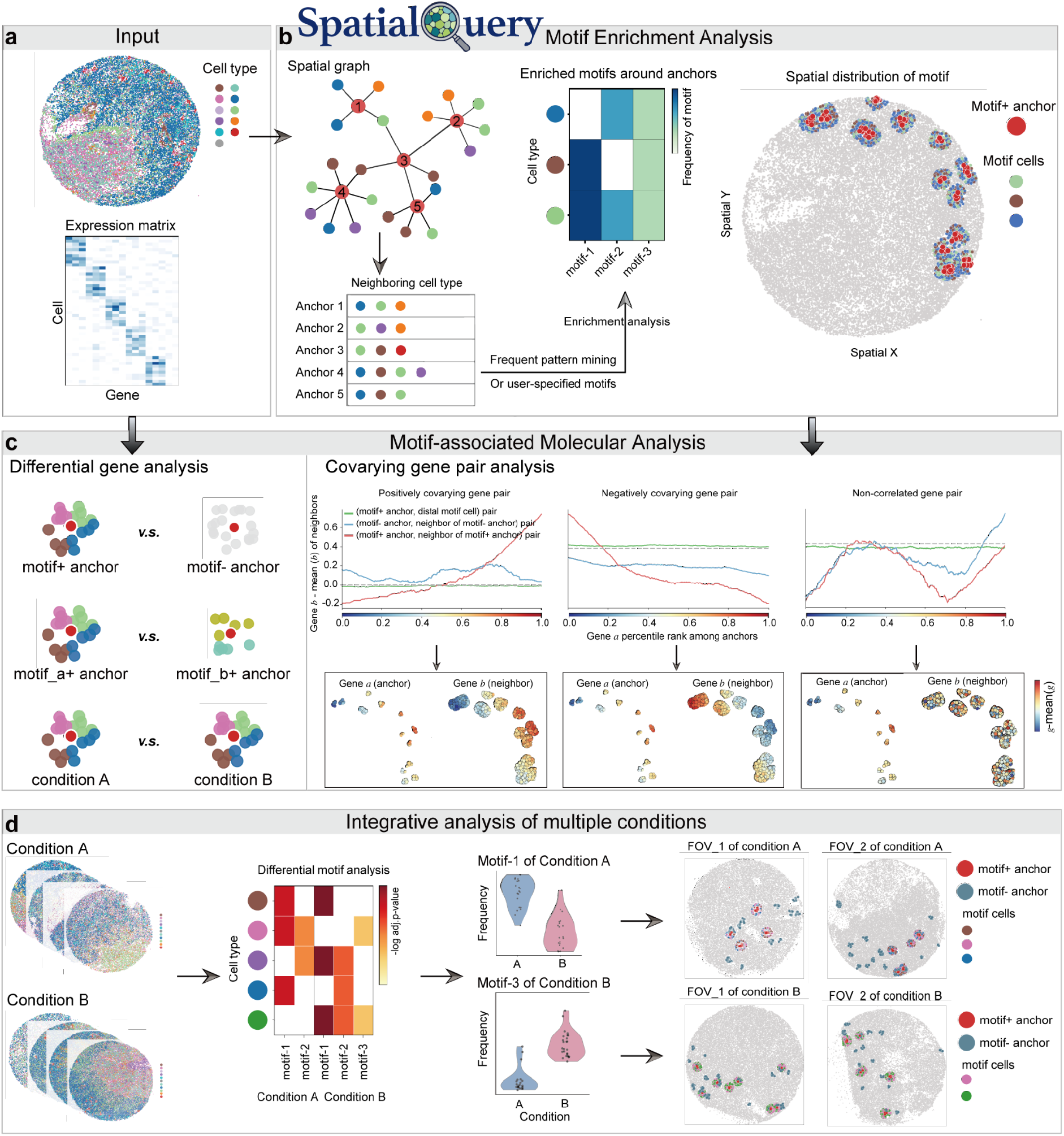
SpatialQuery enables multicellular motif discovery and motif-associated molecular characterization. **a**, SpatialQuery takes spatially resolved data as input, comprising spatial coordinates with cell type annotations and a gene expression or protein abundance matrix. **b**, For motif identification, SpatialQuery constructs a spatial neighbor graph, identifies recurrent cell type combinations via frequent pattern mining or user-specified definitions, and assesses enrichment using a hypergeometric test. **c**, Motif-based molecular analysis. Left, differential gene expression analysis supports three comparisons: anchor cells with versus without a motif in neighborhoods, anchor cells surrounded by different motifs, and anchor cells surrounded by the same motif across conditions. Right, covarying gene pair analysis identifies gene pairs with coordinated expression changes between anchor and neighboring motif cells. Top, anchor gene expression rank (x-axis) versus neighboring cell expression deviation (y-axis) for positively covarying, negatively covarying, and non-correlated gene pairs, respectively. Three lines per panel represent the motif niche (red), non-spatial baseline (green), and non-motif background contexts (blue). Bottom, corresponding spatial expression patterns of gene on anchors (left panel) and paired gene on neighboring motif cells (right panel). **d**, Differential motif analysis identifies condition-specific motifs in multi-condition data, with results shown as a significance heatmap, per-FOV frequency distribution for each motif, and spatial visualization of motif across FOVs.

## Results

### SpatialQuery enables multicellular motif discovery and motif-associated molecular characterization

SpatialQuery takes as input spatial coordinates, cell type annotations, and molecular measurements to identify and characterize cellular motifs (Fig. 1a). We define a cellular motif as a combination of cell types that co-localize around a designated anchor cell type more frequently than expected by chance. SpatialQuery comprises two principal modules: motif discovery to identify statistically enriched motifs, and motif-associated molecular analysis to characterize the transcriptional or proteomic signatures linked to specific cellular motifs.

For motif discovery, the user specifies an anchor cell type. Spatial coordinates are optionally normalized such that one spatial unit corresponds to the mean nearest-neighbor distance, ensuring consistent parameter interpretation across platforms. A k-D tree based spatial graph records the cell type composition of each center cell’s neighborhood. Next, a frequent pattern mining algorithm, FP-Growth^39^, is applied to identify cell type combinations that recurrently surround anchor cells. Hypergeometric testing is used to assess whether observed co-localization frequencies exceed expectations given tissue-wide abundances. Users may restrict analyses to neighborhoods of a minimum size or to motifs exceeding a minimum frequency. After hypergeometric testing, the number of enriched motifs reflects microenvironmental heterogeneity of anchor cells by revealing the number of distinct structural modules. The motif complexity highlights that while some cell types reside in diverse but unstructured environments, others act as organizational hubs. Beyond this unbiased discovery, users can also test whether a specified cell type combination is enriched around a given anchor type (Fig. 1b).

For motif-associated molecular analysis, SpatialQuery provides differential expression analysis and covarying gene pair identification (Fig. 1c). Given an anchor cell type and a motif, three differential expression comparisons are supported: anchor cells with versus without the motif in their neighborhood, anchor cells surrounded by different motifs, and anchor cells within the same motif across experimental conditions. The same comparisons can also be applied to the non-anchor cells in each motif. For covariation analysis, we identify gene pairs exhibiting coordinated expression variation between anchor cells and neighboring motif cells. We compute a shifted Pearson correlation using cell type-specific global means of gene expression as baselines in each field of view (FOV) before testing if the observed correlation is significant against two controls—a non-spatial baseline pairing anchor cells with distal cells of the same cell types as found in the motif, and a non-motif background from anchor cells lacking the motif (see Methods).

For multi-sample studies, neighborhood lists are concatenated across FOVs and enrichment is assessed using a pooled hypergeometric test to identify spatial patterns conserved across biological replicates. When samples are stratified into experimental groups, differential motif analysis is supported by comparing per-FOV motif frequency between groups to uncover condition-specific motifs (Fig. 1d). For atlas-scale studies, an optional scfind-based encoding^40^ compresses sparse expression matrices, substantially reducing memory requirements while preserving essential biological signals^41^.

### SpatialQuery identifies motifs in heterogenous tissues and characterizes gut tube patterning signals

One challenge in cellular motif discovery is that spatially clustered anchor cells may have overlapping neighborhoods, potentially inflating false-positive enrichment signals. To evaluate statistical calibration in a heterogeneous sample, we analyzed a sequential fluorescence in situ hybridization (seqFISH) dataset of mouse embryos at the 8–12 somite stage^35^, where cell types span a wide range of abundances and spatial clustering levels (Fig. 2a). We quantified spatial clustering using Ripley’s L statistic and selected three anchor cell types from distinct levels of elevated spatial autocorrelation - Forebrain/Midbrain/Hindbrain (L = 0.459), Spinal cord (L = 0.287), and Gut tube (L = 0.141) - identified the top motif for each, and recalculated enrichment p-values across 1,000 label permutations. QQ-plots confirmed well-calibrated null p-value distributions across all the three anchor cell types (Supplementary Fig.1).

**Fig. 2.**
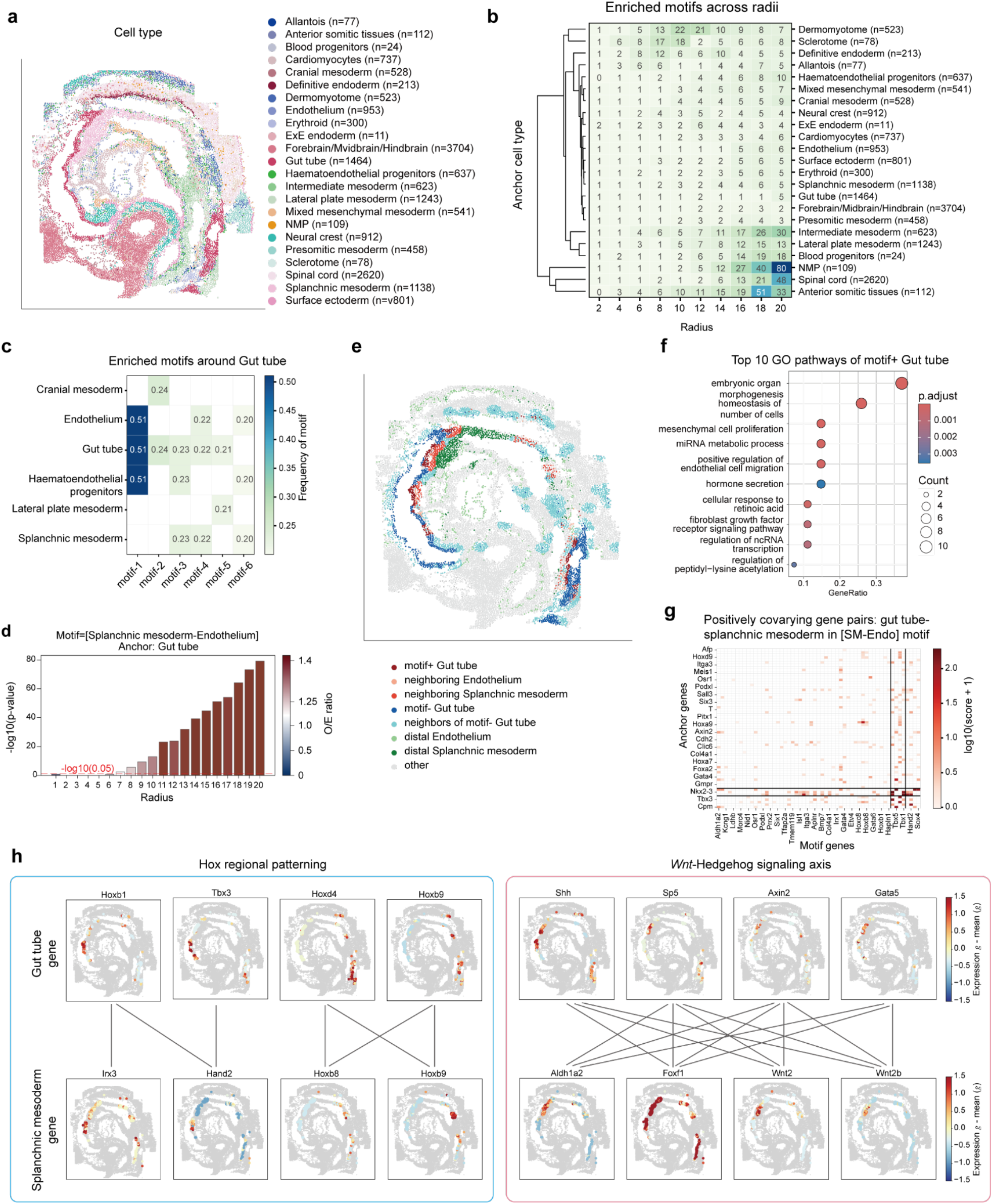
SpatialQuery characterizes patterning signaling of gut tube development. **a**, A mouse embryo at the 8–12 somite stage (seqFISH), colored by cell type with cell counts indicated. **b**, Number of enriched motifs (hypergeometric test, FDR < 0.05) per anchor cell type across neighborhood radii (2–20 spatial units; default frequency threshold = 0.5), with hierarchical clustering of anchor cell types by enrichment profile. **c**, Enriched motifs surrounding gut tube cells (frequency > 0.2, radius=8 spatial units). **d**, Enrichment significance (−log10(*p-*value), y-axis) and observed-to-expected ratio (color) for the [Splanchnic mesoderm–endothelium] motif around gut tube cells across radii. **e**, Spatial distribution of gut tube and motif cells for the [Splanchnic mesoderm-endothelium] motif. **f**, Gene Ontology (GO) enrichment of upregulated genes in motif+ versus motif− gut tube cells. **g**, Positively covarying gene pairs between gut tube (anchor) and splanchnic mesoderm within the [Splanchnic mesoderm–endothelium] motif context. Genes separated by black lines were grouped by biclustering. **h**, Covarying gene pairs between gut tube (top) and splanchnic mesoderm (bottom) within the motif context, organized by Hox regional patterning (left) and Wnt-Hedgehog signaling axis (right). Lines connect covarying gene pairs across the two cell types.

We explored the spatial organization of the mouse embryo by quantifying the number of statistically significant motifs enriched around each anchor cell type across various radii (Fig. 2b). This analysis revealed microenvironmental heterogeneities, wherein some cell types, e.g. the gut tube and Forebrain/Midbrain/Hindbrain, reside in simpler neighborhoods, while others, e.g. anterior somitic tissues, show complex microenvironments. Importantly, there is no direct connection between cell type abundance and neighborhood complexity.

We focused on the gut tube, a central organizer of developmental morphogenesis^42,43^ (Fig. 2c). The identified motifs were biologically coherent: endothelium and haematoendothelial progenitors (motif1, motif6), consistent with the concurrent vascular and hematopoietic specification from common progenitors at E8.5^44,45^; while splanchnic mesoderm and endothelium (motif4), reflect their coordinated roles in gut tube vascularization and mesenchymal patterning^46^. We prioritized the [Splanchnic mesoderm–Endothelium] motif which is a fundamental building block for gut organogenesis. Enrichment analysis confirmed that this spatial association is robust, becoming statistically significant at a radius of 8 spatial units (Fig. 2d). We found 22% of gut tube cells surrounded by this motif within an 8-unit radius and strictly confined to the ventral domain of the gut tube (Fig. 2e), suggesting a position-specific instructive cue. Differential expression analysis comparing gut tube cells surrounded by this motif (Motif+) against those lacking it (Motif-) revealed a ventral foregut identity in Motif+ cells (Supplementary Fig. 2), marked by ventral patterning factors (*Nkx2-3*^46^, *Foxf1*^47^, *Gata2/3/5*^48–50^), and anterior *Hox* genes^42^ (*Hoxa1/Hoxb1, Irx1, Hand1, Osr1, Prrx2*). These cells were also enriched for mediators of epithelial–endothelial and epithelial– mesenchymal crosstalk^51^ (*Podxl* and *Tjp2*), reflecting their physical proximity to the vasculature and mesoderm. Gene Ontology (GO) term analysis confirmed that Motif+ gut tube cells exhibited significant upregulation of pathways critical for organ morphogenesis, endothelial cell migration, and tissue remodeling (Fig. 2f), indicating that this spatial configuration is associated with cell fate specification.

To further dissect the intercellular signaling axes, we quantified gene expression covariation between gut tube and each motif cell type, capturing both positively and negatively coordinated gene programs (Fig. 2g, Supplementary Fig. 3). Unsupervised clustering of the gene-gene correlation matrices revealed three co-regulatory modules, and functional annotation revealed a synchronized developmental program: gut tube genes were enriched for general morphogenesis, mesenchymal development, and epithelial tube morphogenesis. The coupled splanchnic mesoderm covarying genes were enriched for cardiogenesis and vascular support, consistent with the spatial proximity of the ventral foregut to the developing heart field (Supplementary Fig. 4). Parallel analysis of gut tube-endothelium covariation highlighted organ patterning and cardiac vascularization (Supplementary Fig. 4), reinforcing the role of this motif as a multi-lineage coordination hub. From the positively covarying gene pairs between gut tube and splanchnic mesoderm, we identified enrichment of two established developmental programs (Fig. 2h, Supplementary Fig. 5): a HOX patterning module, where anterior and trunk HOX genes (*Hoxb1, Hoxd4, Hoxb9*) covaried with their mesenchymal counterparts (*Hoxb8, Hoxb9, Hand2, Irx3*), reflecting coordinated anterior-posterior regionalization across germ layers; and a *Wnt*-Hedgehog signaling axis, with endodermal *Wnt* targets (*Sp5, Axin2*) and *Shh* coupled to mesenchymal effectors (*Foxf1, Wnt2/2b, Aldh1a2*), consistent with the paracrine loop governing gut morphogenesis.

**Fig. 3.**
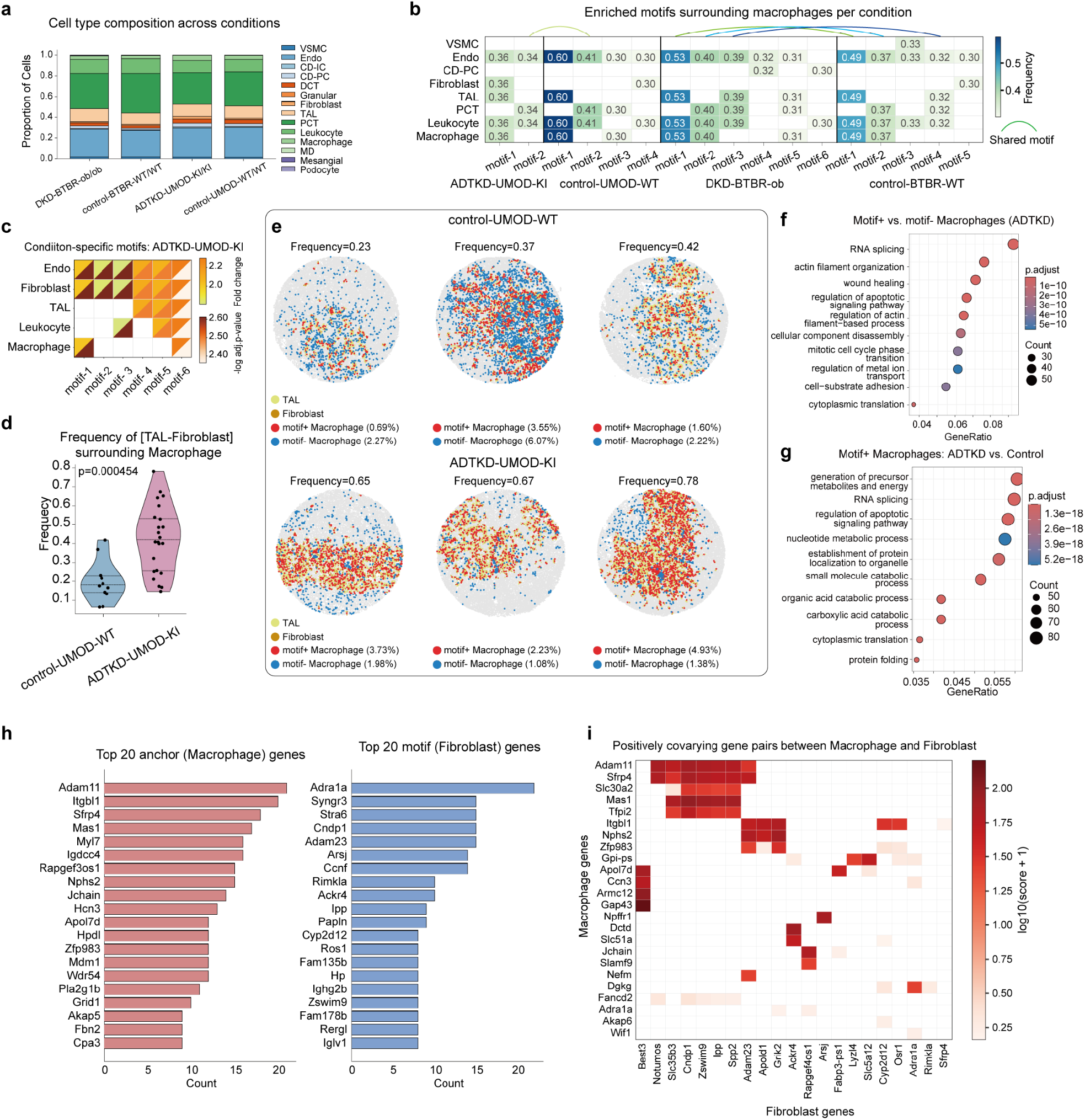
SpatialQuery reveals conserved homeostatic niches and disease-specific fibrotic microenvironments in kidney atlas data. **a**, Cell type composition across four murine kidney conditions. **b**, Enriched motifs surrounding macrophages within each condition. Bracket indicates shared motifs between disease and corresponding control groups. **c**, Enriched motifs of ADTKD-UMOD-KI compared to control-UMOD-WT by differential motif analysis. **d**, Frequency of the [TAL– Fibroblast] motif surrounding macrophages in ADTKD-UMOD-KI versus control-UMOD-WT, quantified per FOV (*p-*value = 4.5 × 10^−^4, Mann-Whitney U test). **e**, Spatial distribution of the [TAL–Fibroblast] motif around macrophages in control-UMOD-WT (top) and ADTKD-UMOD-KI (bottom), showing the three FOVs with highest motif frequency per condition. Percentages indicate the proportion of motif+ or motif-macrophage population within the FOV. **f**, GO enrichment of upregulated pathways in motif+ versus motif-macrophages within ADTKD tissue. **g**, GO enrichment of upregulated pathways in motif+ macrophages from ADTKD versus wild-type control. **h**, Top 20 most frequently appearing anchor (macrophage) and motif (fibroblast) genes among positively covarying gene pairs within the [TAL–Fibroblast] motif context. **i**, Positively covarying gene pairs between macrophage and fibroblast cells within the [TAL–Fibroblast] motif involving top frequent genes.

**Fig. 4.**
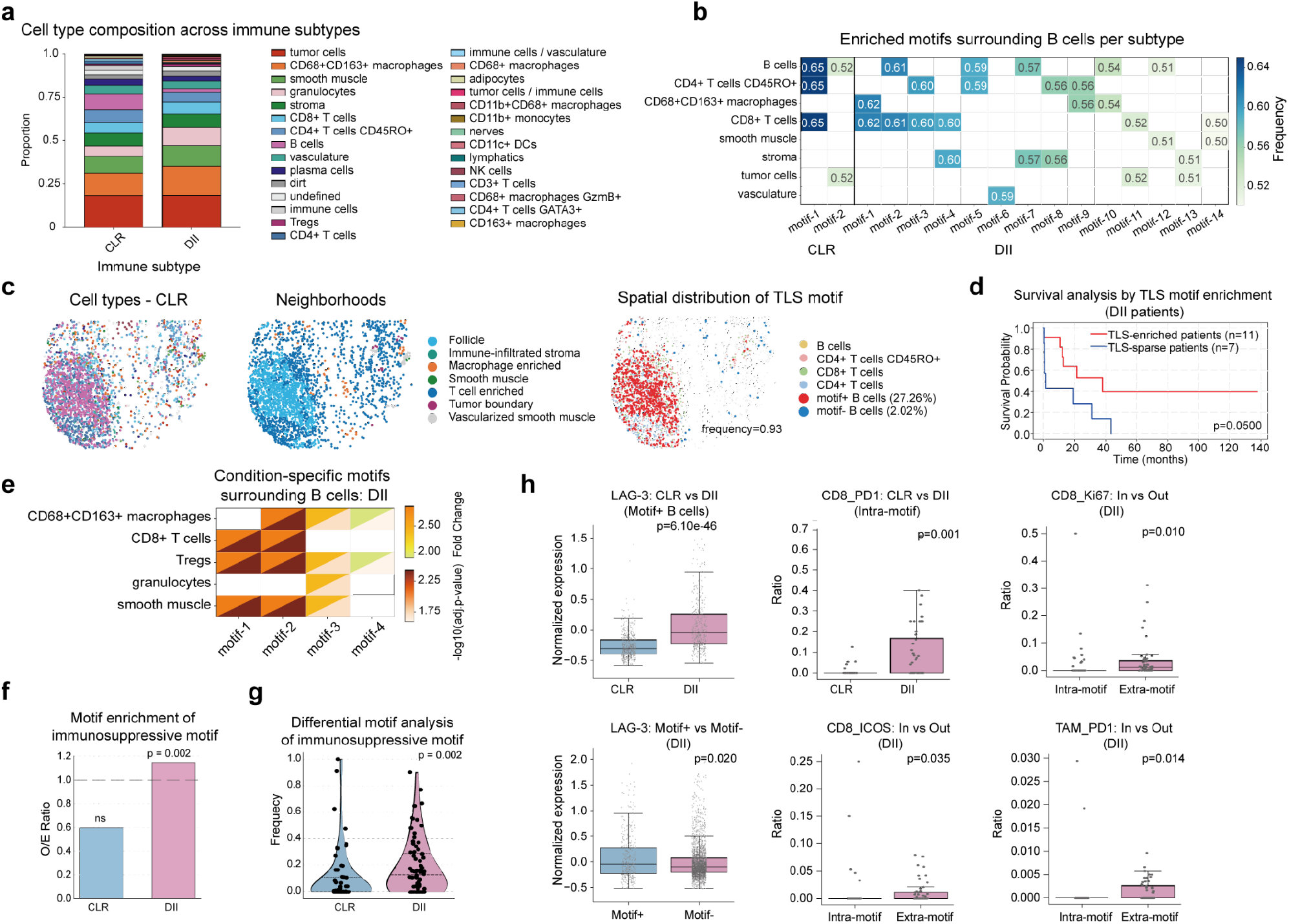
SpatialQuery reveals immunosuppressive microenvironments in colorectal cancer spatial proteomics data. **a**, Cell type composition across Crohn’s-like reaction (CLR, n = 17) and diffuse inflammatory infiltration (DII, n = 18) immune subtypes from CODEX profiling. **b**, Enriched motifs surrounding B cells in CLR and DII patients (radius = 5 spatial units, frequency > 0.5). **c**, Representative CLR FOV showing cell type annotations (left), neighborhood classifications from the original study (middle), and SpatialQuery-identified TLS motif [B cells, CD4+ T cells CD45RO+, CD8+ T cells, CD4+ T cells] distribution (right). **d**, Kaplan-Meier survival analysis of DII patients stratified by TLS motif enrichment. TLS-enriched (≥50% FOVs with significant enrichment, n = 11) versus TLS-sparse (n = 7); *p*-value = 0.05, log-rank test. **e**, DII-specific motifs surrounding B cells compared to CLR. **f**, Observed-to-expected (O/E) ratio for the immunosuppressive motif [CD68+CD163+ macrophages–Tregs–CD8+ T cells–Granulocytes] around B cells, tested independently within CLR and DII patients respectively (hypergeometric test). **g**, Differential motif analysis of the same immunosuppressive motif comparing DII versus CLR patients (*p*-value = 0.002, Mann-Whitney U test). **h**, Differential marker analysis related to the immunosuppressive motif cells. Left column: LAG-3 expression in motif+ B cells comparing CLR versus DII (top; unpaired Mann-Whitney U test of normalized expression data), and motif+ versus motif− B cells within DII (bottom; unpaired Mann-Whitney U test). Middle and right columns: marker positivity ratio in neighboring motif cells of individual FOVs (test with binary values). Middle top, CLR versus DII comparison (unpaired Mann-Whitney U test): PD-1 in CD8+ T cells. Right top and bottom row, intra-motif versus extra-motif within DII (paired Wilcoxon signed-rank test): Ki67 in CD8+ T cells (right top), ICOS in CD8+ T cells (middle bottom), PD-1 in tumor-associated macrophages (TAMs; right bottom).

**Fig. 5.**
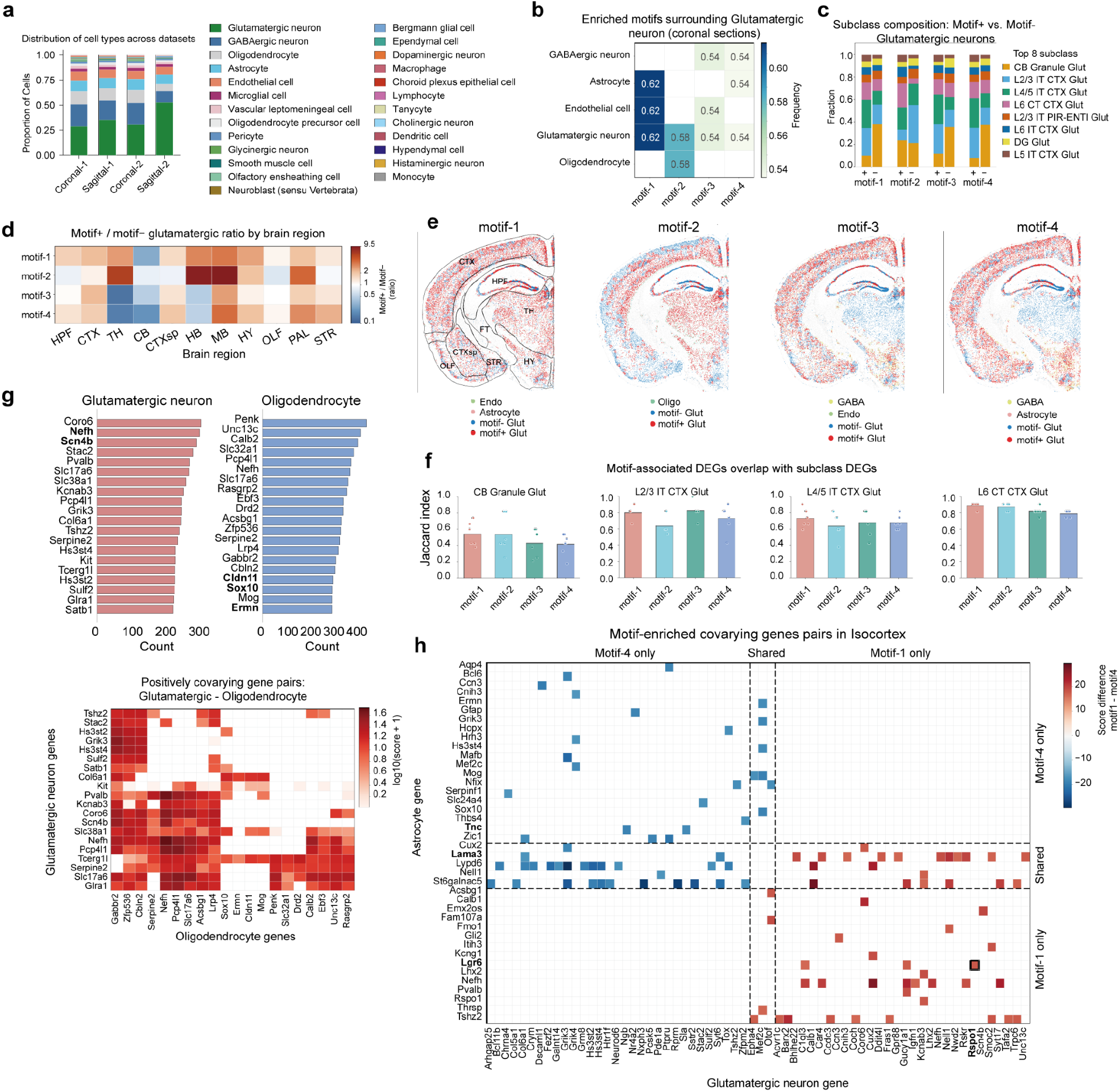
SpatialQuery dissects multicellular organization in a whole-brain MERFISH atlas. **a**, Cell type composition across coronal and sagittal sections from the MERFISH mouse brain atlas (∼8.4 million cells across 239 FOVs). **b**, Enriched motifs surrounding glutamatergic neurons in coronal sections (radius = 5 spatial units, frequency > 0.5). **c**, Subclass composition of motif+ versus motif− glutamatergic neurons for each motif (top 8 subclasses shown). **d**, Ratio of motif+ to motif− glutamatergic neurons in each brain region for each motif. Brain region abbreviations: CTX, isocortex; OLF, olfactory areas; HPF, hippocampal formation; CTXsp, cortical subplate; STR, striatum; PAL, pallidum; TH, thalamus; HY, hypothalamus; MB, midbrain; HB, medulla (hindbrain); CB, cerebellum; FT, fiber tracts; VS, ventricular systems. **e**, Motif-associated cells on a representative coronal section for each motif with major regions outlined. Endo: endothelium; Glut: glutamatergic neuron. **f**, Overlap between motif-context regional DEGs and subclass-level DEGs. For each subclass, bars indicate the mean Jaccard index between the top 20 regional DEGs within each motif context and the corresponding subclass-level top 20 DEGs, averaged across brain regions. Dots represent individual region-level Jaccard values. **g**, Motif-enriched covarying gene pairs between glutamatergic neurons (x-axis) and astrocytes (y-axis), restricted to isocortex. Genes are grouped into motif-4-only, shared, and motif-1-only categories. Bold gene names highlight key genes discussed in the text. **h**, Positively covarying gene pairs between glutamatergic neurons and oligodendrocytes for motif 2 (myelination niche). Left, top 20 most frequently appearing anchor (glutamatergic neurons) and motif (oligodendrocyte) genes among positively covarying gene pairs. Right, positively covarying gene pairs involving the top frequent genes. **i**, Runtime of motif enrichment analysis (left) with whole brain data, and differential motif analysis between coronal and sagittal sections (right) (∼8M cells, 239 FOVs) as a function of neighborhood radius and motif length. Four anchor cell types of varying abundance and 4 motifs of varying lengths are shown.

To assess signaling beyond pairwise interactions in the triad [Gut tube–Splanchnic mesoderm–Endothelium], we compared gene-gene covariation within the three-cell-type motif (N3) against two-cell-type configurations (N2-SM and N2-Endo) (Supplementary Fig. 6). Comparing the difference in covariation scores between N3 and each N2 configuration, we found that N3-enhanced gene pairs were predominantly enriched for morphogen signaling and patterning processes—including *Wnt* pathway, retinoic acid synthesis, and *Hox*-mediated anterior-posterior regionalization—as well as extracellular matrix (ECM) remodeling and vascular guidance. This functional convergence aligns with the established requirement for endoderm-mesoderm-endothelium crosstalk during gut organogenesis^42^. Importantly, these results indicate that the N3 motif captures coordinated signaling programs beyond pairwise analyses.

**Fig. 6.**
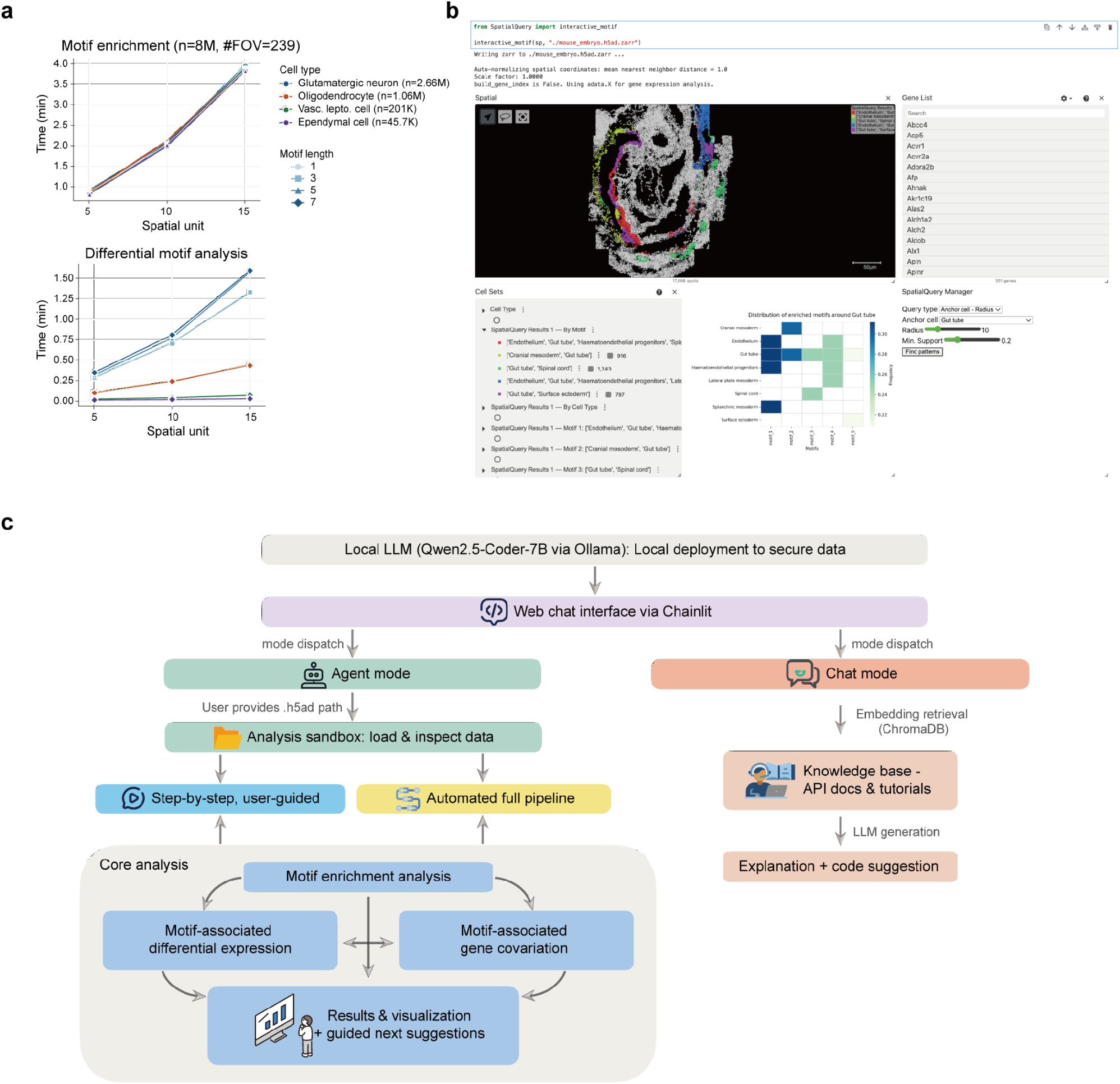
SpatialQuery achieves real-time scalability and supports interactive visual exploration. **a**, Runtime benchmarks on the MERFISH whole-brain atlas. Top, motif enrichment analysis as a function of neighborhood radius for four anchor cell types. Bottom, differential motif analysis between coronal and sagittal sections for four motif lengths. All benchmarks were performed on a MacBook Pro (Apple M2, 8-core CPU, 24 GB RAM). **b**, Integration with Vitessce provides an interactive interface for motif enrichment and visualization. The SpatialQuery Manager panel (bottom right) allows users to specify anchor cell type and parameters. Identified motifs and enrichment statistics are displayed as interactive cell sets (bottom left) and a frequency heatmap (bottom middle). **c**, Architecture of the natural language agent. A locally deployed LLM (Qwen2.5-Coder-7B via Ollama) routes user queries to an agent mode for data loading and pipeline execution, or a chat mode for retrieval-argumented code suggestions and method explanations.

Notably, the signaling axes revealed by SpatialQuery are largely orthogonal to interactions predicted by conventional LR inference tools (Supplementary Fig. 7). Standard LR approaches rely on global co-expression which can mask local heterogeneity, whereas SpatialQuery quantifies motif-associated spatial covariation, detecting coordinated changes. This enables the discovery of spatially restricted regulatory logic that is diluted in tissue-wide averages.

### SpatialQuery reveals conserved homeostatic niches and disease-specific fibrotic microenvironments

To demonstrate SpatialQuery’s capacity to relate spatial organization and changes in gene expression patterns caused by disease, we analyzed a Slide-seqV2 dataset spanning two murine kidney disease models—autosomal dominant tubulointerstitial kidney disease (ADTKD-*Umod*) and diabetic kidney disease (DKD-*ob/ob*)—alongside their respective controls^36^. The integrated dataset comprised approximately 2 million cells in 84 FOVs, annotated into 14 cell types, with broadly similar cell abundances between conditions (Fig. 3a).

Given the pivotal role of macrophages in renal homeostasis and fibrosis^52^, we applied SpatialQuery to characterize macrophage-anchored neighborhoods. Unbiased motif enrichment analysis revealed both conserved and disease-specific patterns (Fig. 3b). Motifs comprising endothelial cells, leukocytes, and proximal or distal tubular segments were consistently detected across disease and control groups, likely representing fundamental anatomical units of the kidney—specifically the juxtaposition of resident macrophages with peritubular capillaries and renal tubules which serves to monitor trans-endothelial transport and maintain tissue homeostasis^53^. Focusing on disease-specific motifs for ADTKD-UMOD we identified six enriched motifs, with [Thick ascending limb (TAL) – Fibroblast – Leukocyte – Endothelial] showing a strong disease association (Fig. 3c). The emergence of fibroblast-containing motifs in ADTKD is consistent with the known pathobiology wherein mutant uromodulin accumulates in TAL cells to trigger interstitial inflammation and progressive fibrosis^54,55^, as well as the established role of macrophage-fibroblast crosstalk in driving fibrosis through reciprocal *Wnt* beta–catenin signaling^56^. We focused subsequent analyses on the [TAL-Fibroblast] motif, which has substantially elevated frequency in ADTKD-UMOD (Fig. 3d,e).

Differential expression analysis comparing Motif+ versus Motif-macrophages in ADTKD revealed upregulation of pathways governing actin filament organization, wound healing, and apoptotic signaling (Fig. 3f, Supplementary Fig. 8), consistent with an activated, remodeling phenotype driven by injury response^57,58^. Furthermore, comparison against motif-associated macrophages in controls revealed that ADTKD motif macrophages are characterized by metabolic reprogramming and protein folding pathways (Fig. 3g). Since ADTKD is fundamentally an endoplasmic reticulum storage disease driven by misfolded uromodulin in TAL cells^54^, the detection of a protein folding signature in neighboring macrophages suggests a potential paracrine response to the proteotoxic stress in neighboring cells, linking the spatial motif directly to the molecular mechanism of the disease^59^.

To dissect the molecular dialogue within the disease-specific [Macrophage-TAL-Fibroblast] motif, we analyzed spatially covarying gene pairs between anchor macrophages and neighboring motif cells (Fig. 3h, Supplementary Fig. 9). We found a pattern of interaction where macrophage genes, including *Adam11, Itgbl1, Sfrp4*, and *Mas1*, showed synchronized expression with specific fibroblast targets. Interestingly, we observed a modular architecture where macrophage genes shared overlapping fibroblast partners, indicating that motif-associated covariation captures coordinated signaling programs (Fig. 3i). For instance, the *Wnt* modulator *Sfrp4* exhibited convergent covariation patterns, indicating a role in driving macrophage-dependent fibrosis^60,61^.

### SpatialQuery reveals immunosuppressive microenvironments in spatial proteomics data

To evaluate SpatialQuery for spatial proteomics data, we analyzed a CODEX dataset profiling the colorectal cancer invasive front, comprising Crohn’s-like reaction (CLR, 17 patients) and diffuse inflammatory infiltration (DII, 18 patients) subtypes^37^. Each patient contributed four tissue microarray cores, yielding 140 FOVs with ∼258k cells (Fig. 4a). We explored anchor cell microenvironments with a neighborhood radius of 5 spatial units. Since the original study focused on tertiary lymphoid structures (TLSs), lymphoid aggregates scaffolded by a fibroblast network which are important for organizing anti-cancer immunity^62^, we focused our analysis on B cells. Unbiased motif enrichment analysis revealed a structural divergence between subtypes. In CLR patients, B cells were frequently embedded within immune-rich neighborhoods containing CD4+ T cells CD45RO+ and CD8+ T cells, forming early-stage TLSs. In contrast, DII motifs were heterogeneous, including M2-polarized tumor-associated macrophages (CD68+CD163+ macrophages, TAMs), stroma, vasculature, and tumor cells alongside lymphocyte populations (Fig. 4b), suggesting a more immunosuppressive microenvironment.

To demonstrate SpatialQuery’s capacity for supervised queries, we specified a TLS motif as [B cell-CD4+ T cell–CD8+ T cell–CD4+ T cell CD45RO+]. Although this motif was enriched in both patient groups, the enrichment magnitude differed substantially and consistently across multiple neighborhood radii (Supplementary Fig. 10a). In CLR patients, 31.4% of B cells (3,225/10,269) were surrounded by this TLS motif, compared to only 9.0% (249/2,774) in DII patients (p<0.0001 by Fisher’s exact test, radius = 5 spatial units). In CLR patients, 94.8% of TLS motif+ cells localized to T cell-enriched or Follicle neighborhoods defined in the original study, compared to 68.7% for DII patients (p<0.0001 with Fisher’s exact test, Supplementary Fig. 10b). We also found elevated localization to Macrophage-enriched (11.6% vs. 1.8%, p<0.0001) and Immune-infiltrated stroma regions (9.0% vs. 0.4%, p<0.0001). This spatial “mislocalization” indicates that even when B-T cell aggregates occur in DII patients, they are embedded within immunosuppressive rather than lymphoid-supportive contexts (Fig. 4c). Stratification of DII patients based on TLS motif enrichment prevalence across their FOVs (see Methods) followed by Kaplan-Meier survival analysis revealed improved overall survival for TLS-enriched patients (log-rank test, p=0.05; Fig. 4d), demonstrating that spatial motifs can be directly linked to clinical outcomes.

Differential motif analysis revealed that B cells in DII patients are surrounded by motifs containing TAMs, CD8+ T cells, Tregs, and granulocytes, suggesting an immunosuppressive microenvironment in DII patients (Fig. 4e). We defined an immunosuppressive motif as [CD68+CD163+ macrophage–Treg–CD8+ T cell–granulocyte], and found enrichment around B cells in DII (p=0.002), but not in CLR patients (Fig. 4f-g). To characterize the functional state of immune cells within the immunosuppressive microenvironment, we compared the proportion of cells expressing each marker within versus outside the DII-specific motif cells, as well as between CLR and DII patients within motifs (Mann-Whitney U test; Fig. 4h). For anchor cells, motif+ B cells in DII patients showed elevated expression of the exhaustion marker *LAG-3* relative to both motif-B cells of DII patients (p=0.020) and motif+ B cells in CLR patients (p<0.0001)^63^. For non-anchor cells within the motifs, CD8+ T cells showed a terminally exhausted phenotype characterized by higher proportion of PD-1 in DII compared to CLR patients (p=0.001). They also showed lower Ki67 (p=0.010), and ICOS (p=0.035) relative to CD8+ T cells found outside of motifs in DII patients. Interestingly, intra-motif TAMs not only displayed lower PD-1 positivity than extra-motif TAMs (p=0.014), but the pattern of PD-1+Ki67−ICOS− CD8+ T cells coexisting with PD-1− TAMs deviates from classical T cell exhaustion and the proliferative response observed following successful PD-1 blockade^64^. Since PD-1− TAMs may induce immunosuppression through checkpoint-independent mechanisms such as TGF-β and IL-10 secretion^65–68^, these spatial motifs may represent microenvironments where CD8+ T cell dysfunction is sustained, reducing the efficacy of anti-PD-1 therapy^69,70^.

### SpatialQuery reveals the interplay between regional identity and motif-associated programs in a whole-brain MERFISH atlas

To evaluate SpatialQuery in a tissue with complex spatial architecture, we applied it to a whole-brain MERFISH dataset comprising ∼8.4 million murine cells profiled with a 1,122-gene panel across 239 FOVs (Fig. 5a)^38^. We designated glutamatergic neurons, which play key roles in sending excitatory signals and regulating brain blood flow^71^, as anchor cells. Applying SpatialQuery to 213 coronal sections, we identified four significantly enriched spatial motifs around glutamatergic neurons (Fig. 5b, radius=5 spatial units, motif frequency > 0.5). Motifs 1 and 3 both contain endothelial cells, suggesting neurovascular unit (NVU) organizations. While motif 1 [glutamatergic-astrocyte-endothelial] likely reflects canonical activity-dependent blood flow regulation^72^, motif 3 incorporates GABAergic interneurons, consistent with bidirectional modulation of vascular tone through vasoactive neuropeptides^73^. Motif 4 [glutamatergic-astrocyte-GABAergic] reflects the tripartite synapse^74,75^ and motif 2 [glutamatergic-oligodendrocyte] likely captures activity-dependent myelination^76,77^. Notably, while motifs 1, 3, and 4 involve cell types associated with neuronal signaling related to real-time physiological regulation, motif 2 involves cell types implicated in structural maturation governing long-term axonal integrity.

Comparison of subclass compositions between Motif+ and Motif-glutamatergic neurons showed that those found in motifs 1, 3, and 4 are dominated by cortical subclasses (L2/3 IT, L4/5 IT, L6 CT, L6 IT), with cerebellar granule glut substantially depleted (Fig. 5c). This is consistent with the distinct cytoarchitecture of the neocortex, where layer-specific glutamatergic neurons are densely intermixed with protoplasmic astrocytes and local GABAergic interneurons, while granule cells are common in the cerebral cortex^78^. Regional enrichment analysis corroborated this pattern: motifs 1, 3, and 4-positive glutamatergic neurons are depleted in cerebellum (Fig. 5d-e). Motif 1 shows broad enrichment across most regions, consistent with the ubiquitous requirement for neurovascular coupling^72^, whereas motifs 3 and 4 are preferentially enriched in midbrain, pallidum, and striatum–regions with dense GABAergic circuitry^79^. Motif 2 is most enriched in cerebellum, midbrain, and hindbrain, white-matter-rich regions where activity-dependent myelination demand is high^80,81^.

To investigate if the motif context shapes transcriptomic identity, we performed differential expression analysis across brain regions while controlling for subclass composition. The top region-enriched genes were highly consistent across motifs, and clustering patterns of regions were preserved across motifs (Supplementary Fig. 11-14). Notably, these motif+ regional DEG profiles closely recapitulated those of corresponding subclasses, regardless of motif context (Fig. 5f)^38,82,83^. That is, the transcriptomic profiles of glutamatergic neurons are predominantly driven by subclass and regional identity rather than by the microenvironment (Supplementary Fig. 15-16).

We investigated the interplay between large-scale regional organization and multicellular microenvironments by carrying out gene-gene covariation analysis between the anchor and other motif cells. Since gene expression in glutamatergic neurons is driven primarily by region rather than motif, we hypothesized that the same would be true for the covarying programs. Indeed, across all four motifs, the top covarying gene pairs were dominated by region-associated markers, (e,g., subcortical markers in the glutamatergic neurons, *Zic1*^84^ and *Zic4*^85^, and *Slc17a6*^38^, cortical markers *Slc17a7*^38^, and region-restricted astrocyte genes, *Agt*^86^ and *Gabbr2*^87^) (Supplementary Fig. 17, 18a-c). Interestingly, motif 2 [glutamatergic neuron–oligodendrocyte] displayed a globally conserved axis of gene-gene covariation. We observed synchronized expression between markers of high-caliber projection neurons^88^ (*Nefh****/****Scn4b*) and the oligodendrocyte myelination machinery^89–91^ (*Cldn11, Sox10, Ermn*) across the entire brain (Fig. 5g). Such motif-associated covariation stems from the large number of genes involved in myelination^91^, which provides sufficient signal-to-noise ratio to overcome regional variance, while it’s less prominent in the more transient signaling pathways characterizing the other three motifs.

To further examine the microenvironmental effects, we directly compared the covarying gene pairs for glutamatergic neuron–astrocyte, between motif 1 (NVU) and motif 4 (GABAergic– astrocyte niche). We selected the top 100 motif-specific covarying gene pairs from each context (Supplementary Fig. 18d). Despite the fact that there were no pairs shared between the two contexts, the individual genes involved showed notable overlap. This could be explained by the fact that the intercellular communication of neurovascular coupling and neurotransmitter cycling operate predominantly through post-translational mechanisms, with the motif-specific transcriptional signatures being less important^92^. Taken together, we conclude that variation in gene expression is primarily driven by regional identity rather than motif context^93^.

Next, we tested whether motif-associated covariation signals could nonetheless be recovered after controlling for regional heterogeneity. We restricted the [glutamatergic– astrocyte] analysis to isocortex (Supplementary Fig. 17), and repeated the comparison between motif 1 and motif 4. Region stratification reduced gene overlap between the two motif contexts (Fig. 5h, Jaccard index decreased from 0.392 to 0.047 for center genes and from 0.361 to 0.125 for surrounding genes), and revealed distinct motif-associated signals (Supplementary Fig. 19). Specifically, motif 1 revealed preferential covariation involving astrocyte-expressed *Lama3*, which is enriched in perivascular astrocytes and implicated in NVU basement membrane architecture^94,95^. We also identified the glutamatergic neuron-to-astrocyte *Rspo1*-*Lgr6* LR pair, consistent with *RSpo1*-mediated Wnt/beta-catenin signaling through *LGR6* receptors on cortical layer 2/3 and 5 astrocytes^96^. In contrast, the motif 4 [glutamatergic-GABAergic–astrocyte] was enriched for covariation involving astrocyte-expressed *Tnc*, implicated in perisynaptic ECM remodeling at inhibitory synapses^97^. These region-stratified results confirm that SpatialQuery can uncover motif-associated covariation programs that are masked by regional identity variance at the whole-brain level^98,99^. Collectively, these findings highlight SpatialQuery’s capacity to identify biologically relevant multicellular programs through spatial covariation. While robust programs are detectable brain-wide, subtler programs may require region-stratified analysis to disentangle motif-specific signals from regional heterogeneity.

### SpatialQuery achieves real-time scalability and supports interactive web based deployment

SpatialQuery achieves computational efficiency through efficient data structures and closed-form statistical tests, enabling real-time exploratory analysis at atlas scale. We benchmarked three key analyses, frequent pattern mining, motif enrichment analysis, and differential motif analysis, on both a single sample and three atlas datasets (with between 18K and 8M total cells) across varying settings (Fig. 6a, Supplementary Fig. 20-21). Motif enrichment analysis completed within seconds on single-FOV data, and took <10 minutes for atlas data for typical parameter configurations (neighborhood size ranging from 5 to 10). Differential motif analysis across all 239 FOVs took <2 minutes even for neighborhood radius=15. Runtime scaled linearly with neighborhood radius and was largely independent of motif length.

We have taken steps to move SpatialQuery beyond the current paradigm of computational tools which are downloaded and run locally. Thanks to its computational efficiency, SpatialQuery can be deployed as a web service, and as a proof of concept it is being integrated into the HuBMAP data portal^100^ through Vitessce^101^ (Fig. 6b). Users are provided an interactive interface where they can specify anchor cell types and analysis parameters, and explore the resulting motifs and enrichment statistics directly. For users running SpatialQuery locally, we provide a natural language agent that supports end-to-end analysis through conversational queries (Fig. 6c). The agent operates on a locally deployed language model so that no user data leaves the local machine, and offers two interaction modes: an agent mode that loads spatial omics data and executes the analytical workflow without the need for writing python code, and a chat mode that provides method explanations and code suggestions via retrieval-augmented generation from SpatialQuery’s API documentation (Supplementary Fig. 22).

## Discussion

We have introduced SpatialQuery, a computational framework that unifies multicellular spatial motif discovery with motif-associated molecular characterization from spatially resolved single-cell data. By connecting spatial pattern identification directly to downstream differential expression and cross-cell covariation analysis, SpatialQuery enables a coherent analytical workflow from motif discovery to molecular interpretation. Applications across four diverse biological systems and spatial platforms illustrate this capacity.

Compared to existing methods for exploring spatial domains, SpatialQuery offers several new perspectives. Cellular motifs can be identified either in a supervised or unsupervised manner, and the ability to focus on an anchor cell type is broadly useful as researchers often have existing hypotheses about which cell type is most relevant for their study. The use of cellular motifs means that SpatialQuery is capable of discovering tissue heterogeneities unlike most other neighborhood finding methods which implicitly assume tissue homogeneity. Moreover, SpatialQuery can investigate multi-cell interactions, something that would be prohibitive if applied to all possible combinations of three or more cell types. The cross-cell covariation module quantifies coordinated deviations from the global baseline between anchor and neighboring motif cells rather than co-expression. This design captures signals beyond known ligand-receptor interactions, including shared microenvironmental responses and synchronized transcriptional programs.

Nevertheless, SpatialQuery still has several limitations. The identified gene pairs are correlational rather than causal: observed associations may reflect direct intercellular signaling, common upstream regulators, or residual confounders such as sub-regional variation in cell type composition not fully controlled by the analytical design. Moreover, covariation analysis computes a population-level statistic across all motif-positive cells, implicitly assuming transcriptional coherence within the group. In tissues with strong tissue-level compartmentalization, region-driven expression differences can dominate the covariation signal and obscure motif-specific programs. Although stratified analysis can disentangle these sources of variance, as demonstrated in the mouse brain analysis, compartment annotations are not always available and the two signals may be conflated.

SpatialQuery bridges motif discovery and motif-associated molecular characterization, two analytical tasks that have largely been addressed by separate tools in the spatial omics field. As such, it provides a useful tool as researchers seek to identify functional tissue units^102^ to understand how groups of cells cooperate to support tissue functionality. With the increasing availability of atlas-scale spatial omics datasets, its real-time efficiency, flexible query-driven design and scalability to millions of cells position it as a practical framework for systematic, hypothesis driven, dissection of how multicellular spatial organization shapes cellular function.

## Methods

### SpatialQuery workflow

SpatialQuery is a computational framework for identifying recurrent multicellular spatial patterns, i.e., motifs, and characterizing their associated molecular signatures from spatially resolved single-cell data. The framework operates on three input data types: two-dimensional spatial coordinates representing cellular positions within tissue sections, cell type or cell state annotations, and gene expression matrices from spatial transcriptomics or protein abundance measurements from spatial proteomics platforms. Following data preprocessing, the analytical pipeline proceeds through two principal modules: frequent cellular motif discovery with statistical enrichment assessment, and motif-associated molecular analysis, including differential expression analysis and coordinated gene-pair identification.

#### Data preprocessing and spatial indexing

To ensure consistent parameter interpretation across varying imaging resolutions and platforms, SpatialQuery provides optional coordinate normalization. When enabled, spatial coordinates are divided by the mean nearest-neighbor distance computed per FOV, such that one spatial unit corresponds to one mean cell spacing. This normalization ensures that neighborhood parameters, such as radius thresholds, carry equivalent biological meaning across heterogeneous datasets. When the native coordinate system already encodes a physically meaningful unit (e.g., micrometers), users may bypass normalization and operate on the original coordinates.

A k-dimensional tree (k-D tree) is subsequently constructed from the two-dimensional coordinates to facilitate efficient spatial neighborhood queries. The k-D tree is built through recursive median-based binary partitioning, alternating between two spatial coordinates at successive tree levels. This construction procedure achieves *O*(*n log n*) time complexity for *n* input points and produces a balanced tree structure with uniform distribution of spatial points. The balanced architecture ensures *O*(*log n*) expected time complexity for nearest-neighbor and range queries, enabling scalable analysis of datasets comprising hundreds of thousands to millions of cells.

For gene expression data, SpatialQuery provides two processing pathways tailored to different data modalities. For spatial transcriptomics data, the framework applies standard log-normalization as in Scanpy^103^. For spatial proteomics data, where marker abundances often follow distinct distributional properties, feature-wise z-score transformation is applied to standardize measurements across markers. Alternatively, expression data can be encoded using scfind^40^, a cell atlas indexing algorithm that compresses sparse expression matrices through quantile-based discretization while preserving efficient query capabilities. This encoding substantially reduces memory requirements for large-scale datasets and enables rapid binary expression queries. Analyses performed on scfind-indexed data operate on binarized expression states, which has been shown to preserve essential biological signals for cell type identification and pattern discovery while improving computational tractability.

#### Frequent cellular motif discovery

The motif discovery module identifies cell type combinations exhibiting recurrent spatial co-localization relative to a specified anchor cell type. For each cell of the designated anchor type, a spatial neighborhood is defined using either k-nearest neighbors (kNN) or a fixed radius threshold, with neighbors retrieved via k-D tree queries. The cell type composition of each neighborhood is recorded as a transaction—a set containing the cell type labels of all neighboring cells. This formulation transforms the spatial pattern discovery problem into a frequent itemset mining problem. The Frequent Pattern Growth (FP-Growth) algorithm^39^ is employed to identify cell type combinations occurring above a certain fraction (minimum frequency support) of all anchor cell neighborhoods. Importantly, the FP-Growth algorithm achieves linear time complexity with respect to the number of frequent patterns. Since any subset of a frequent pattern is necessarily also frequent, the algorithm may identify many redundant patterns. To eliminate redundant output, only maximal frequent patterns— patterns possessing no frequent proper superset at the specified support threshold—are retained. These maximal patterns, termed cellular motifs, represent the most complete cell type combinations that recurrently surround the anchor cell type filtered by frequency threshold.

The minimum support parameter is set by the users, and it controls the trade-off between motif length and discovery sensitivity. At a given threshold, FP-Growth identifies maximal frequent patterns. Lower frequency thresholds permit longer motifs (containing more cell types within each motif), because the support requirement is relaxed. However, longer motifs are not necessarily statistically significant by hypergeometric test (see Motif enrichment analysis), as the number of neighborhoods containing the full combination may be small. In such cases, shorter subsets of a non-significant maximal pattern may themselves be significant. SpatialQuery reports maximal patterns from FP-Growth together with hypergeometric p-values, allowing users to identify patterns and sub-patterns based on cardinality, enrichment, and statistical significance. We recommend exploring multiple frequency thresholds and neighborhood sizes to characterize the motif landscape at different levels of complexity, or specify the cell type combinations of interest as motifs to explore.

#### Motif enrichment analysis

While the FP-Growth algorithm identifies motifs that frequently co-localize with a given anchor cell type, high frequency alone does not establish statistical significance. A motif may appear frequently around an anchor type simply because the constituent cell types are abundant in the tissue. To assess whether observed co-localization exceeds random expectation, a hypergeometric test is applied. For a candidate motif M and anchor cell type C, the hypergeometric distribution models the probability of observing *k* or more anchor cells with motif M in their neighborhood, given: the total number of cells *N* in the tissue, the number of neighborhoods (across all cell types) containing motif M, and the total number of anchor cells of type C. The one-sided *p*-value is computed as *P*(*X* ≥ *k*), representing the probability of observing the given or greater enrichment under the null hypothesis that anchor cells are randomly distributed among all neighborhoods containing the motif. When multiple motifs are tested simultaneously, *p*-values are adjusted for multiple hypothesis testing using the Benjamini-Hochberg procedure to control the false discovery rate at a specified threshold (default *α* = 0.05).

#### Motif-associated differential gene expression analysis

To characterize the molecular signatures associated with motifs, we provide differential expression analysis comparing cells within versus outside motif neighborhoods. Given an anchor cell type C and motif M, three biologically relevant representative comparisons are supported:

1. Anchor cells whose neighborhoods contain the motif (motif+ anchors) are compared with cells of the same type whose neighborhoods lack the motif (motif-anchors). This comparison identifies genes in the anchor cell type whose expression is associated with the presence of the specific neighboring cell type combination, potentially reflecting niche-dependent transcriptional programs.
2. Anchor cells surrounded by different motifs (M1+ anchors versus M2+ anchors) are compared, revealing genes associated with distinct microenvironmental compositions.
3. Anchor cells with identical motif neighborhoods but from different experimental conditions or disease states are compared, enabling identification of condition-specific gene expression changes within defined spatial niches.

Statistical testing employs methods appropriate to the expression data representation. For continuous expression data stored in the original matrix format, differential expression testing is performed using either the Wilcoxon rank-sum test or Welch’s t-test as in Scanpy. For scfind-indexed expression data, Fisher’s exact test is applied to 2×2 contingency tables with binary data comparing expression frequencies between cell groups. Genes are prefiltered to require expression in at least a minimum fraction of cells (default 5%) in at least one comparison group, and resulting *p*-values are adjusted using the Benjamini-Hochberg procedure.

#### Motif-specific co-varying gene pairs identification

##### Cell partition

We provide functionality to discover gene pairs whose cross-correlation between anchor cells and neighboring motif cells is elevated in the spatial context. For a given anchor cell type C and motif M, cells are separated into distinct groups based on their spatial relationships. Anchor cells are divided into those whose neighborhoods contain the motif (M+ anchors) and those of the same type whose neighborhoods lack the motif (M− anchors). Similarly, motif-constituent cells are classified into those spatially neighboring C+ anchors (Mn) and distal cells (Mnn). We also define the general neighborhood of motif-negative anchors (Nnm), comprising all direct neighbors of M-anchors regardless of cell type. Using these partitions, we construct three comparative spatial contexts:

1. The motif niche (M+ anchor, Mn) formed by spatial pairing between anchor cells and their corresponding motif neighbors, representing the spatial context of motif M.
2. The non-spatial baseline (M+ anchor, Mnn) is constructed by synthetically pairing M+ anchors with distal Mnn cells, which randomizes spatial proximity while retaining cell type identities.
3. The non-motif background (M-anchor, Nnm) formed by spatial pairings between M-anchors and their local neighbors, which controls for non-specific spatial autocorrelation and the baseline microenvironmental state of the anchor cell type when the specific motif is absent. When the motif comprises multiple cell types, the analysis can be computed either by pooling all motif cell types into a single population, or separately for each constituent cell type within the motif, depending on the biological question of interest.

##### Shifted Pearson correlation

To quantify gene co-variation patterns across distinct spatial contexts while preserving signals relative to tissue-wide baselines, we implement a global-mean-centered Pearson correlation metric. Let *ci* denote the cell type identity of cell *i*. For any gene *g*, we define the reference baseline 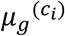 as the global mean expression of gene *g* computed across all cells of type *c*_*i*_ in the FOV. For a defined set of cell pairs *P* = {(*i, j*)} determined by the above three spatial contexts, the shifted cross-cell correlation between gene *g* in cell type *ci* and gene *h* in cell type *cj* is computed as:

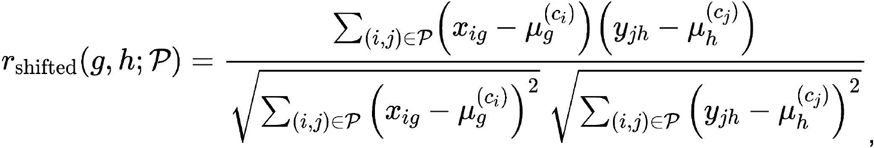

where *x*_*ig*_ and *y*_*ih*_ denote the normalized expression levels. Correspondingly, the three shifted cross-cell correlation matrices for each gene pair are computed as 1) *r*_*n*_ which quantifies relationships within the motif niche (M+ anchor, M_n_), 2) *r*_*nn*_ using all-to-all pairs in spatial context (M+ anchor, M_nn_), and 3) *r*_*nm*_ for non-motif background (M-anchor, N_nm_), respectively.

##### Statistical testing

To test whether the shifted Pearson correlation for a given gene pair differs significantly between two spatial contexts, we apply Fisher’s Z-transformation to convert each correlation coefficient *r* to an approximately normally distributed score:

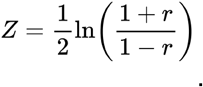

Given two transformed scores *Z*_1_ and *Z*_2_ derived from contexts with effective sample sizes *n*_1_ and *n*_2_, the difference is evaluated using the test statistic

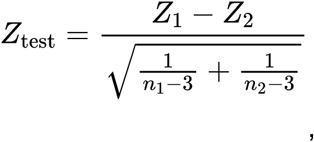

which follows a standard normal distribution under the null hypothesis that the two underlying correlations are equal. To account for spatial autocorrelation and the dependency structure inherent in cell-pair data, we conservatively define the effective sample size as *n*_*eff*_ = *min*(*n*_*center*_, *n*_*neighbor*_). We perform two independent two-tailed hypothesis tests for each gene pair:

1. *r*_*n*_ against *r*_*nn*_: testing whether spatial proximity to anchor cells specifically alters gene-gene correlation compared to the same cell types located distally.
2. *r*_*n*_ against *r*_*nm*_: testing whether the correlation is specific to the motif context compared to anchor cells lacking the motif.

Both tests detect gene pairs with either significantly evaluated or reduced correlations, as both positive and negative covariation patterns may carry biological significance. Candidate gene pairs are required to satisfy directional consistency: for each test, we define the correlation difference *Δr* = *r*_*n*_ − *r*_*control*_, where *r*_*control*_ is the corresponding correlation from the comparison context (*r*_*nn*_ for test 1, *r*_*nm*_ for test 2). The sign of *Δr* must agree across both tests, filtering out incoherent signals where spatial and context drivers exert opposing effects. P-values from directionally consistent pairs were pooled and adjusted for multiple hypothesis testing using the Benjamini-Hochberg procedure.

##### Scoring and ranking

Finally, we prioritize gene pairs using a combined score (*S*_*combined*_) that integrates effect sizes and significance levels from both statistical comparisons:

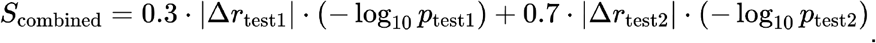

We apply a higher weight to the second test to prioritize signals where the anchor cell state explicitly differs depending on the presence of the specific spatial motif, thereby highlighting context-driven intercellular coordination. Gene pairs are ranked by the absolute value of the combined score.

##### Comparison of covariation across contexts

To identify gene pairs whose covariation is preferentially associated with a specific context, we compare covariation results for the same anchor-motif cell type pair between two analytical scenarios. These scenarios may represent distinct motif contexts surrounding the same anchor cell type, or the same motif context under different experimental conditions. For each scenario, covariation analysis was performed independently as described above, yielding a combined score for each gene pair. Gene pairs that were statistically significant (adjusted p-value < 0.05) in at least one scenario, or that appeared among the significant pairs in both scenarios, are selected as candidates. For each candidate gene pair, the difference in combined scores between the two scenarios is computed. Gene pairs are then ranked by this difference, and those in the top and bottom user-specified percentiles are designated as context-enriched pairs for each respective scenario. This procedure was applied to compare glutamatergic neuron-astrocyte covariation between motif 1 (NVU) and motif 4 (GABAergic-astrocyte niche) within isocortex (Fig. 5h), and to compare the three-cell-type motif (N3) against two-cell-type configurations (N2) surrounding gut tube cells (Supplementary Fig. 6).

#### Multi-sample integrative analysis

##### Cohort-level aggregation and motif discovery

To enable cross-sample comparison, individual FOVs are stratified into user-defined conditions. Preprocessing and neighboring graph construction are performed independently for each FOV as described previously. For condition-level motif discovery, neighbor transaction lists are aggregated across all constituent FOVs prior to FP-Growth pattern mining. Similarly, global enrichment significance is assessed using a pooled hypergeometric test, where the contingency table parameters—total cell count, anchor cell count, and motif-positive neighborhood count—are summed across all FOVs within the condition.

##### Differential motif analysis

For comparisons between two conditions, candidate motifs are first identified by extracting maximal frequent patterns (above a user-defined support frequency threshold) from each FOV across both conditions. For each candidate motif, the motif frequency—defined as the proportion of anchor cells whose neighborhoods contain the motif—is computed independently within each FOV, yielding a distribution of frequency values per condition. The two-sided Mann-Whitney U test is then applied to assess whether motif frequencies differ significantly between conditions, with Benjamini-Hochberg correction for multiple testings. Alternatively, the user can specify a motif of interest and differential testing is performed directly on the specified motif.

#### Binary-based molecular analysis with scfind index

To enable scalable analysis of atlas-level datasets, SpatialQuery provides an optional expression encoding scheme based on scfind^40^. When operating on scfind-indexed data, differential expression analysis employs Fisher’s exact test on 2 × 2 contingency tables comparing the frequency of expressing versus non-expressing cells between two groups. For covariation analysis, gene expression states are represented as binary (expressing/non-expressing) and the shifted correlation is computed on these binary vectors using the same framework as continuous data. Analyses performed on scfind-indexed data operate on binarized expression states, which has been shown to preserve essential biological signals for cell type identification and pattern discovery while improving computational tractability (Supplementary Fig. 23-24).

### Calibration of p-values

We assessed whether SpatialQuery maintains statistical calibration under varying degrees of spatial autocorrelation by performing null-simulation analysis. We first quantified the magnitude of spatial clustering for each cell type using Ripley’s L function (*squidpy*.*gr*.*ripley*). Three anchor cell types spanning different clustering levels were selected: Forebrain/Midbrain/Hindbrain (high clustering, mean L=0.459), Spinal cord (L = 0.287), and Gut tube (intermediate, mean L = 0.141). For each anchor type, we identified the top-ranked motif using a search radius of 8 spatial units and a frequency threshold of 0.5. We then generated 1,000 null datasets by randomly permuting cell type labels while preserving spatial coordinates. This procedure disrupts true cell type-spatial relationships while maintaining overall tissue architecture. We then calculated enrichment p-values of the top-ranked motif for each permuted dataset using the hypergeometric test. Well-calibrated tests should produce p-values following a Uniform(0,1) distribution under the null hypothesis; we evaluated this using quantile-quantile plots comparing observed to theoretical uniform quantiles.

### Comparison with ligand-receptor inference

To assess the relationship between SpatialQuery covariation signals and conventional ligand-receptor predictions, we applied CellChatV2^23^, COMMOT^28^, CellPhoneDB^24^, and Scotia^27^ to the mouse embryo dataset using the CellChatDB LR database with default parameters. Predicted interactions between gut tube and splanchnic mesoderm/endothelium were extracted and compared with positively covarying gene pairs identified by SpatialQuery within the [splanchnic mesoderm-endothelium] motif context.

### TLS motif stratification and survival analysis

We stratified DII patients based on TLS motif prevalence, with the TLS motif defined as the combination of B cells, CD4+ T cells, CD8+ T cells, and CD4+ T cells CD45RO+. For each patient, TLS motif enrichment was tested independently within each FOV using the hypergeometric test. Patients with significant TLS enrichment (FDR < 0.05) in at least 50% of their FOVs were classified as TLS-enriched (n = 11); the remaining patients were classified as TLS-sparse (n = 7). Overall survival was compared between the two groups using Kaplan-Meier estimation and the log-rank test. TLS motif+ cell proportions between CLR and DII patients were evaluated using Fisher’s exact test. Overlap between TLS motif-positive B cells and cellular neighborhoods defined in the original study was quantified per condition, and differences in localization to specific neighborhood categories were assessed using Fisher’s exact test.

### Dataset processing

#### Mouse embryo seqFISH

Spatial transcriptomics data of mouse embryos at the 8–12 somite stage^35^ profiles 17,806 cells across using a 351-gene seqFISH panel. Cell type annotations were taken from the original study. Spatial coordinates were provided in 0.1 micrometer/pixel and were normalized by mean nearest-neighbor distance. Gene expression data were log-normalized using Scanpy (scanpy.pp.normalize_total followed by scanpy.pp.log1p).

#### Mouse kidney Slide-seqV2

High-resolution Slide-seqV2 spatial transcriptomics data spanning two murine kidney disease models^36^. The dataset comprises approximately 2 million cells and 84 FOVs across four conditions: ADTKD-UMOD knock-in (ADTKD-UMOD-KI), wild-type control (UMOD-WT), diabetic kidney disease (DKD-BTBR-ob/ob), and wild-type control (BTBR-WT). Cell type annotations were taken from the original study. Spatial coordinates in micrometers were normalized by mean nearest-neighbor distance. For covariation analysis, FOVs within each condition were concatenated, and the top 3,000 highly variable genes were identified from the concatenated data following log-normalization by Scanpy. Gene-gene shifted correlations were then computed on these genes.

#### Colorectal cancer CODEX

CODEX spatial proteomics data profiling the colorectal cancer invasive front^37^ comprises 35 patients classified into Crohn’s-like reaction (CLR, n = 17) and diffuse inflammatory infiltration (DII, n = 18) subtypes, with each patient contributing four tissue microarray cores for a total of 140 FOVs and approximately 258,000 cells. A panel of 56 markers was used for cell phenotyping; cell type annotations and cellular neighborhood assignments were taken from the original study. Spatial coordinates in micrometers were normalized by mean nearest-neighbor distance. Feature-wise z-score transformation is applied to standardize measurements across markers.

#### MERFISH mouse brain atlas

Whole-brain MERFISH data comprises approximately 8.4 million cells profiled with a 1,122-gene panel across 239 tissue sections (213 coronal, 26 sagittal) from four adult mice^38^. Cell type annotations at the class, subclass, and supertype levels, as well as brain region assignments, were taken from the original study. Brain region annotations were used for region-stratified analyses. Spatial coordinates in micrometers were normalized by mean nearest-neighbor distance. Normalized expression data from the original study was used for molecular analysis.

### Software implementation

SpatialQuery is implemented in Python 3.10+ and operates on AnnData objects for compatibility with the Scanpy (V1.11.5) single-cell analysis ecosystem. Spatial indexing utilizes scipy.spatial.KDTree with multi-threaded query execution (SciPy V1.15.3). FP-Growth implementation is provided by the mlxtend library (V0.23.4).

### Scalability benchmarking

All benchmarks were performed on a MacBook Pro with Apple M2 chip (8-core CPU) and 24 GB RAM running macOS Tahoe V26.2. Reported times represent single-run wall-clock time for computation only, excluding data loading.

## Data availability

The mouse embryo seqFISH dataset is available from https://crukci.shinyapps.io/SpatialMouseAtlas/. The kidney Slide-seqV2 dataset is available from CELLxGENE database (https://cellxgene.cziscience.com/collections/8e880741-bf9a-4c8e-9227-934204631d2a). The colorectal cancer CODEX dataset is available from Mendeley(https://dx.doi.org/10.17632/mpjzbtfgfr.1). The MERFISH whole-brain atlas data are available from CELLxGENE database (https://cellxgene.cziscience.com/collections/0cca8620-8dee-45d0-aef5-23f032a5cf09).

## Code availability

SpatialQuery is available as an open-source Python package at GitHub repository (https://github.com/ShaokunAn/Spatial-Query) under the MIT license. Documentation is available at https://spatialquery.readthedocs.io/en/latest/. The natural language Agent model for end-to-end analysis is at GitHub repository (https://github.com/ShaokunAn/SpatialQuery-Agent). Analysis scripts for reproducing the results are available at https://spatialquery.readthedocs.io/en/latest/tutorials/index.html. A ready-to-use analysis template is available through HuBMAP workspaces (https://portal.hubmapconsortium.org/templates/spatial_query; registration required), where users can run SpatialQuery analyses directly on HuBMAP-hosted datasets in a pre-configured environment.

## Acknowledgements

We thank Jingyi Cao and Jae-Won Cho for helpful discussions and the rest of the Hemberg lab for constructive feedback on the manuscript. We thank Alexa Guan, Yufei Wang, and Joshua Choi for critical evaluation of biological analyses. This work was supported by the NIH (3OT2OD033759-01S5, U24 CA268108, OT2 OD033758).

## Author contributions

M.H. conceived the study. S.A. and M.H. designed the analytical framework. S.A. and M.K. implemented the software. S.A. performed all analyses. S.A. and M.H. wrote the manuscript with contributions from M.K. and N.G. All authors reviewed and approved the final manuscript.

