## supplementary figures for "SpatialQuery: scalable discovery and molecular characterization of multicellular motifs from spatial omics data"

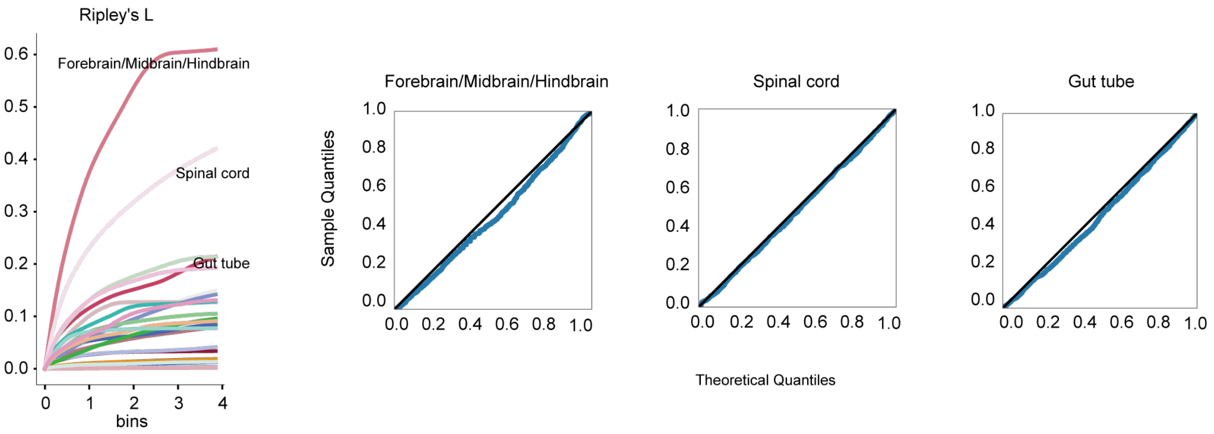


Supplementary Fig. 1 | Calibration of p-values under varying spatial autocorrelation. Left, Ripley's L statistic for all cell types in the mouse embryo dataset, plotted across evaluation distances (bins), with three representative anchor cell types spanning different clustering levels highlighted: Forebrain/Midbrain/Hindbrain (L = 0.459), Spinal cord (L = 0.287), and Gut tube (L = 0.141). Right, QQ-plots of null p-value distributions for motif enrichment (hypergeometric test) from 1,000 label permutations for each anchor cell type. For each anchor type, the top-ranked motif was identified with original data, cell type labels were randomly permuted while preserving spatial coordinates, and enrichment p-values were recalculated of the same motif. Black diagonal indicates the theoretical Uniform(0,1) distribution. Radius = 8 spatial units, frequency threshold = 0.5.

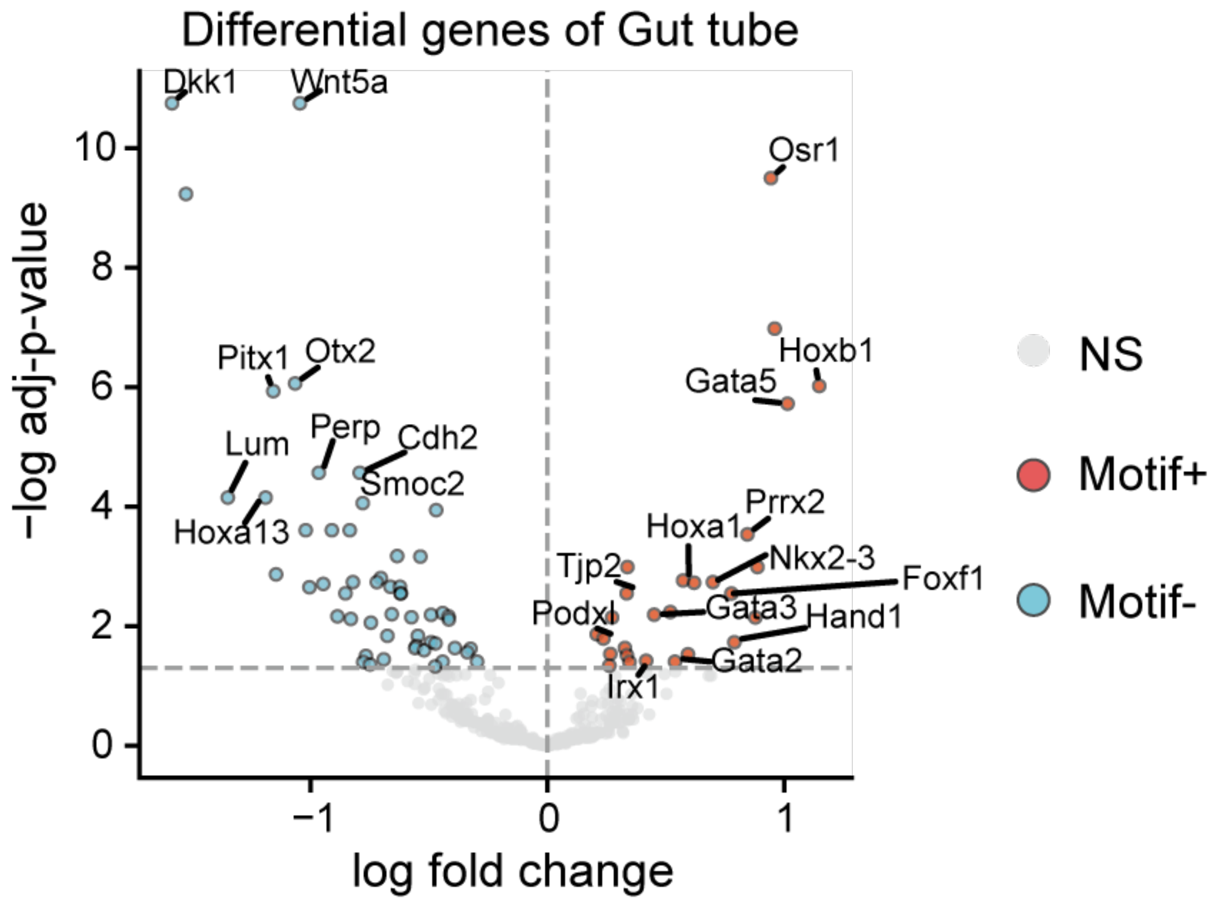


Supplementary Fig. 2 | Differential gene expression between motif+ and motif− gut tube cells. Volcano plot showing differentially expressed genes between gut tube cells with versus without the [Splanchnic mesoderm–Endothelium] motif in their neighborhood. Genes significantly upregulated in Motif+ (red) and Motif− (blue) gut tube cells are highlighted. Radius = 8 spatial units.

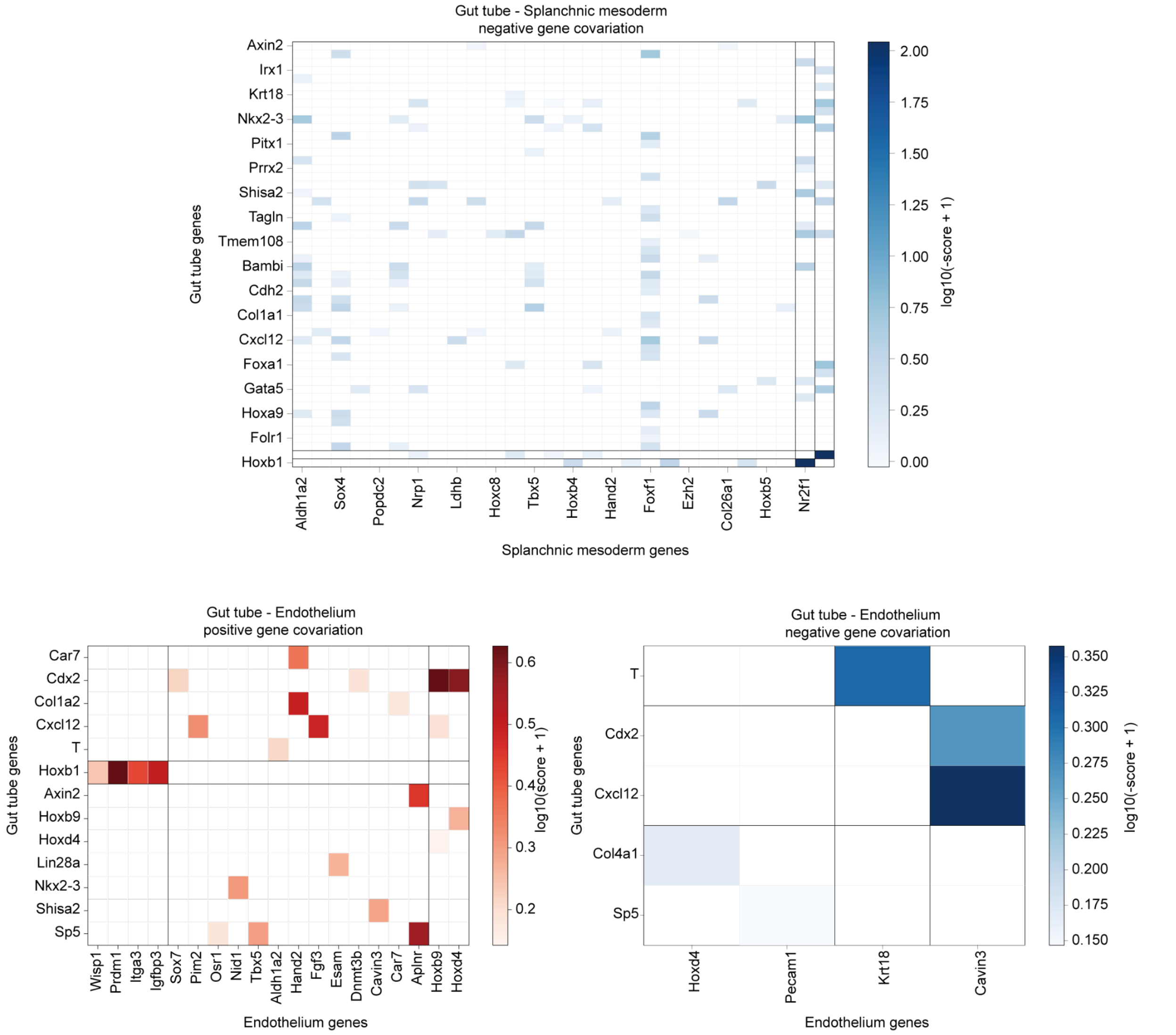


Supplementary Fig. 3 | Covarying gene pairs between gut tube and motif cell types. Top, negatively covarying gene pairs between gut tube and splanchnic mesoderm. Bottom left, positively covarying gene pairs between gut tube and endothelium. Bottom right, negatively covarying gene pairs between gut tube and endothelium. Rows (gut tube genes) and columns (motif genes) are organized by biclustering into three row clusters and three column clusters. All analyses performed within the [Splanchnic mesoderm–Endothelium] motif context. Genes separated by black lines were grouped by biclustering.

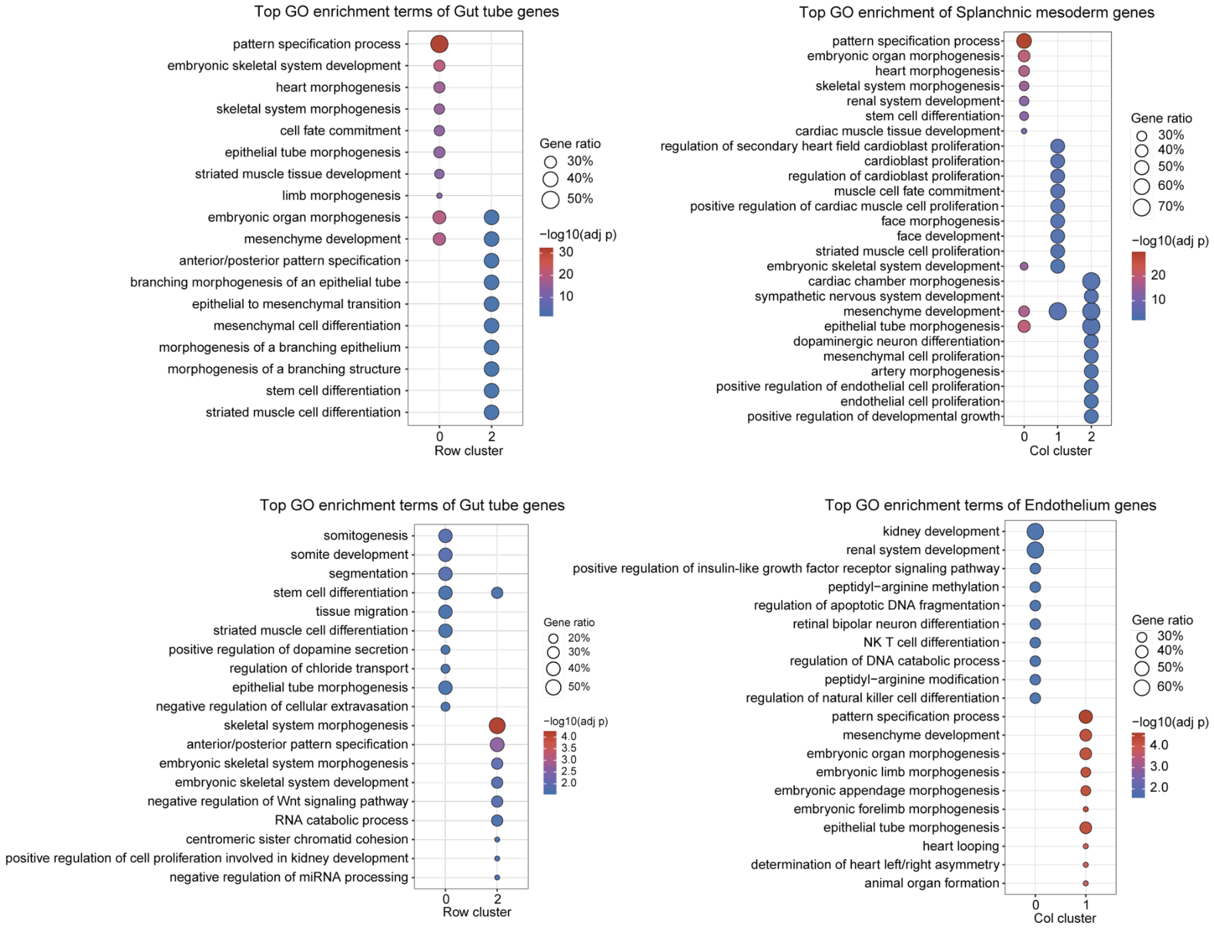


Supplementary Fig. 4 | GO enrichment of genes involved in covarying gene pairs. Top row, GO enrichment of gut tube genes (left) and splanchnic mesoderm genes (right) from gut tube–splanchnic mesoderm covarying gene pairs, with biclustered covarying gene pair modules (Fig. 2g) that are grouped by row and column cluster respectively. Bottom row, GO enrichment of gut tube genes (left) and endothelium genes (right) from gut tube–endothelium covarying gene pairs, grouped by cluster. All analyses performed within the [Splanchnic mesoderm–Endothelium] motif context.

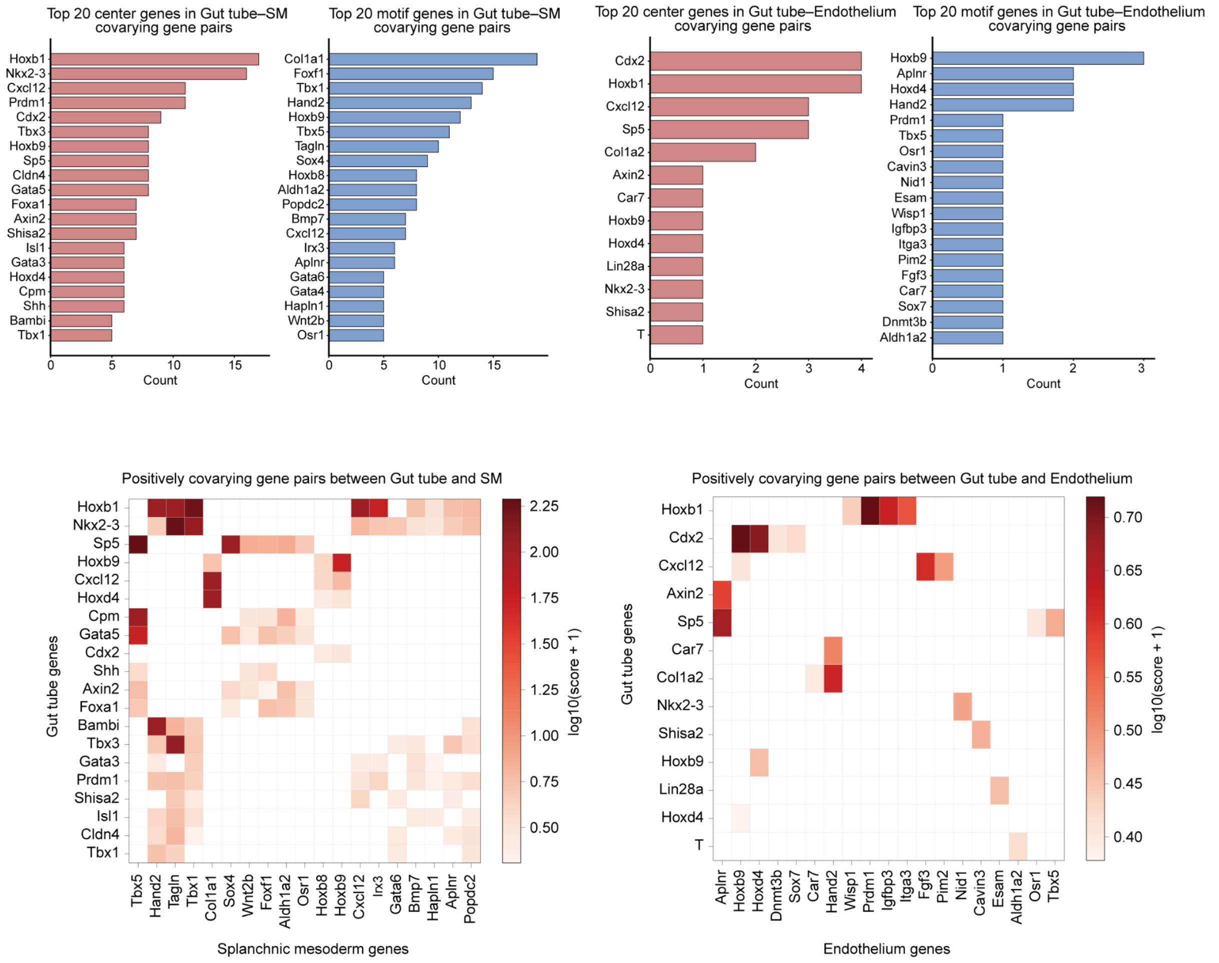


Supplementary Fig. 5 | Top frequent genes in covarying gene pairs between gut tube and motif cell types. Top, distribution of top 20 anchor (gut tube, red) and motif genes appearing in positively covarying gene pairs for gut tube–splanchnic mesoderm (left) and gut tube–endothelium (right) with [Splanchnic mesoderm–Endothelium] motif context. Bottom, corresponding covarying gene pair matrices involving these top frequent genes.

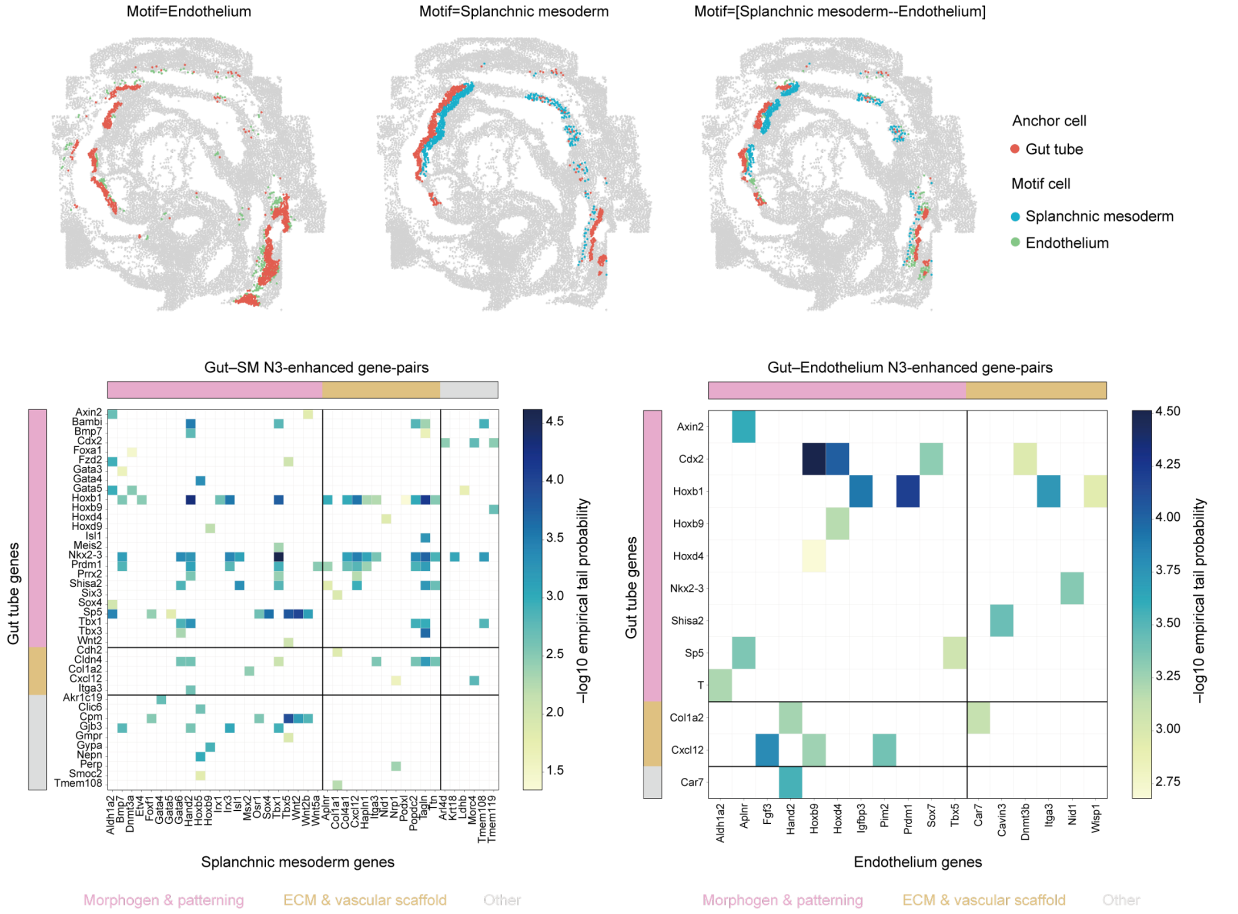


Supplementary Fig. 6 | Comparison of multi-cellular and pairwise neighborhood analyses. Top, spatial distribution of gut tube anchor cells and motif cells under pairwise configurations (motif = Endothelium only; motif = Splanchnic mesoderm only) versus the three-cell-type motif [Splanchnic mesoderm, Endothelium]. Bottom, gene pairs with enhanced covariation in the three-cell-type context (N3) relative to pairwise configurations (N2), shown for gut tube–splanchnic mesoderm (left) and gut tube–endothelium (right). Gene categories are annotated along the axes: morphogen and patterning (magenta), ECM and vascular scaffold (teal).

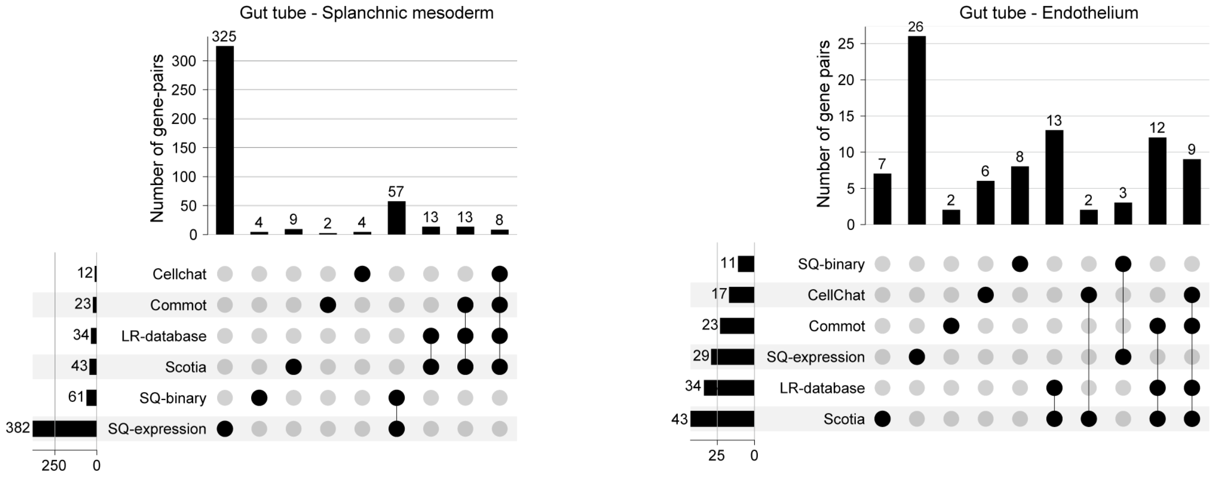


Supplementary Fig. 7 | Comparison with ligand-receptor interaction tools. UpSet plots showing overlap between covarying gene pairs identified by SpatialQuery (expression-based and binary-based) within [Splanchnic mesoderm–Endothelium] motif context, and gene pairs predicted by CellChat, COMMOT, Scotia, and curated LR databases, for gut tube–splanchnic mesoderm (left) and gut tube–endothelium (right).

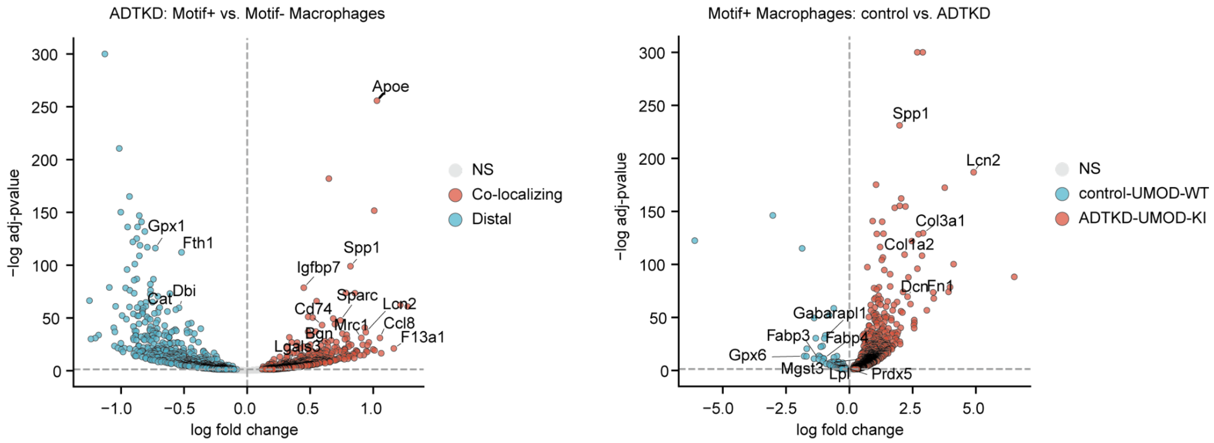


Supplementary Fig. 8 | Differential gene expression analysis of macrophages in kidney disease. Left, volcano plot comparing Motif+ versus Motif- macrophages within ADTKD tissue for motif [TAL–Fibroblast]. Right, volcano plot of differentially expressed genes in motif+ macrophages between ADTKD-UMOD-KI and control-UMOD-WT.

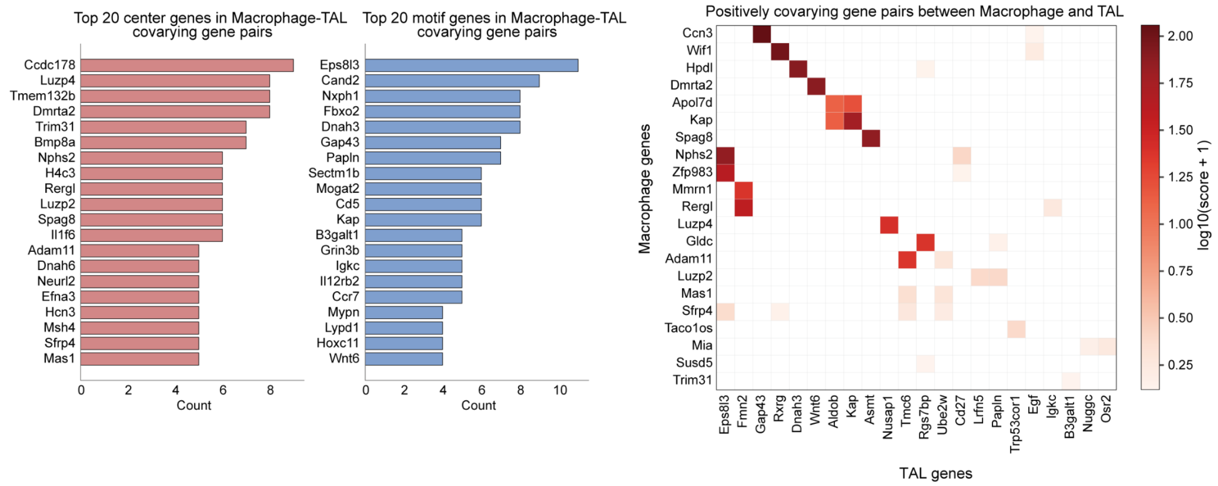


Supplementary Fig. 9 | Covarying gene pairs between macrophage and TAL within the [TAL–Fibroblast] motif. Left, distribution of the top 20 anchor (macrophage, red) and motif (TAL, blue) genes among positively covarying gene pairs. Right, covarying gene pair matrix between macrophage and TAL cells involving top frequent genes.

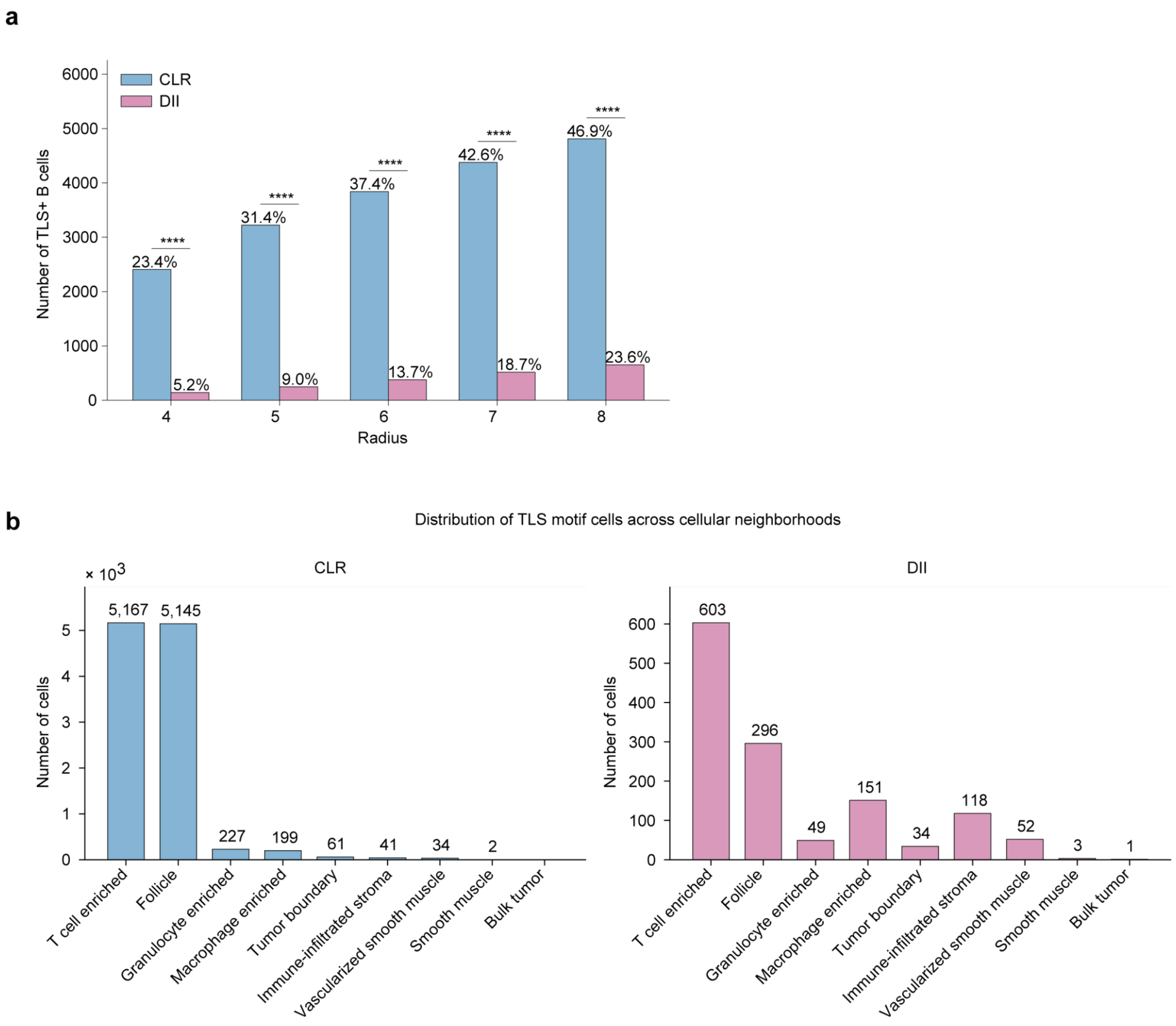


Supplementary Fig. 10 | TLS motif abundance in CLR and DII cohorts. a, Number of TLS motif+ B cells (motif = [B cells, CD4+ T cells, CD8+ T cells, CD4+ T cells CD45RO+]) across neighborhood radii from 4 to 8 spatial units in CLR and DII patients. Percentages indicate the proportion of B cells surrounded by this motif. Fisher's exact test; ****, $p < 0.0001$. b, Distribution of TLS motif-associated cells across cellular neighborhood categories defined in the original study, for CLR (left) and DII (right). Radius = 5.

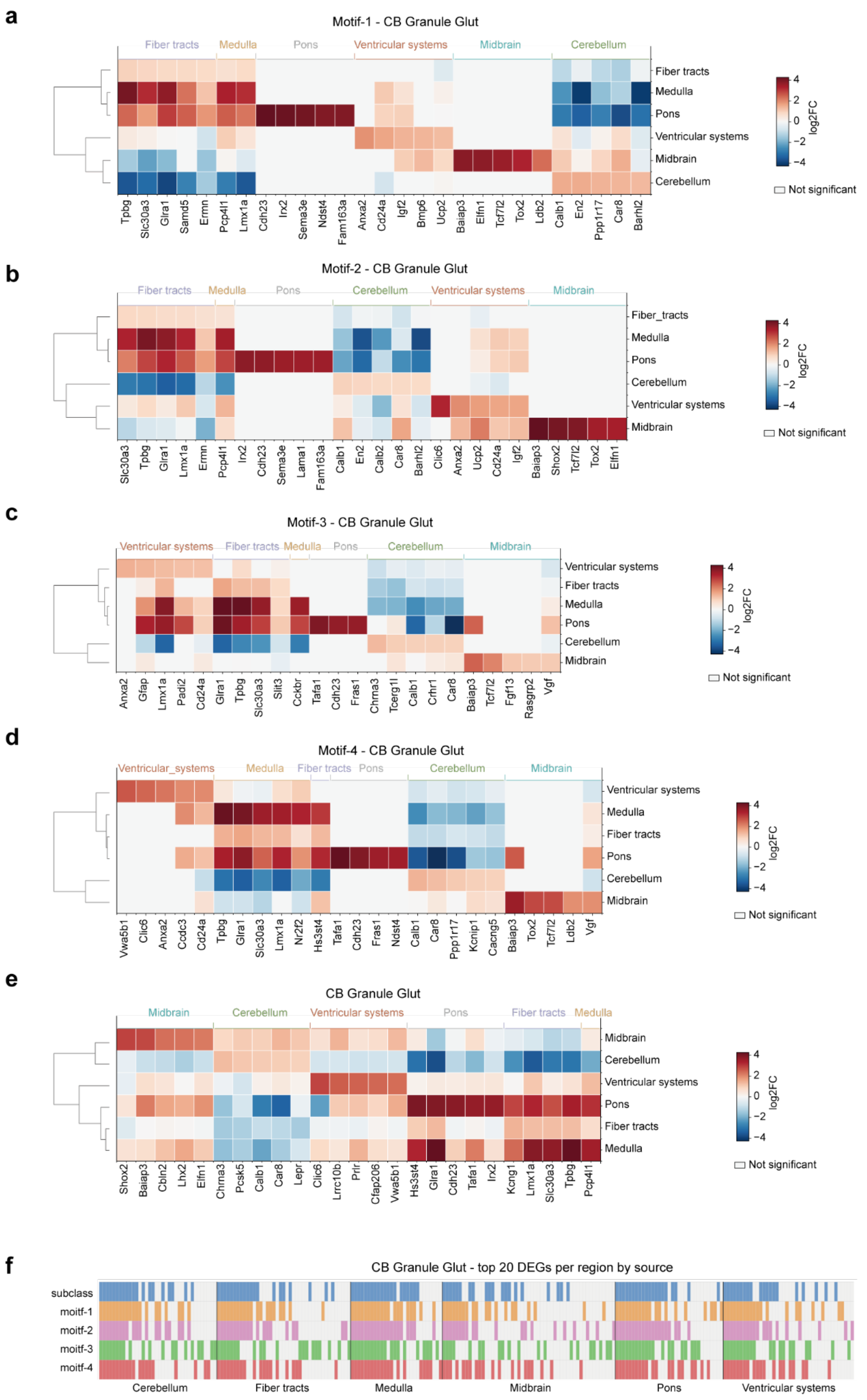


Supplementary Fig. 11 | Regional differential expression of motif+ glutamatergic neurons: CB Granule Glut subclass. a–d, Top 5 differentially expressed genes per brain region (row) for motif 1 (a), motif 2 (b), motif 3 (c), and motif 4 (d), controlling for subclass by restricting to CB Granule Glut neurons. Hierarchical clustering of regions (dendrogram) was performed on enrichment scores of individual genes (columns). e, Subclass-level regional DEGs without motif stratification. Log2FC values reflect expression in each region versus remaining regions. f, Categorical heatmap showing the union of motif-context and subclass-level regional DEGs per region. Each column represents a gene. Colored entries indicate that the gene ranks among the top 20 regional DEGs for the corresponding row source; gray entries indicate absence from that source's top 20 list.

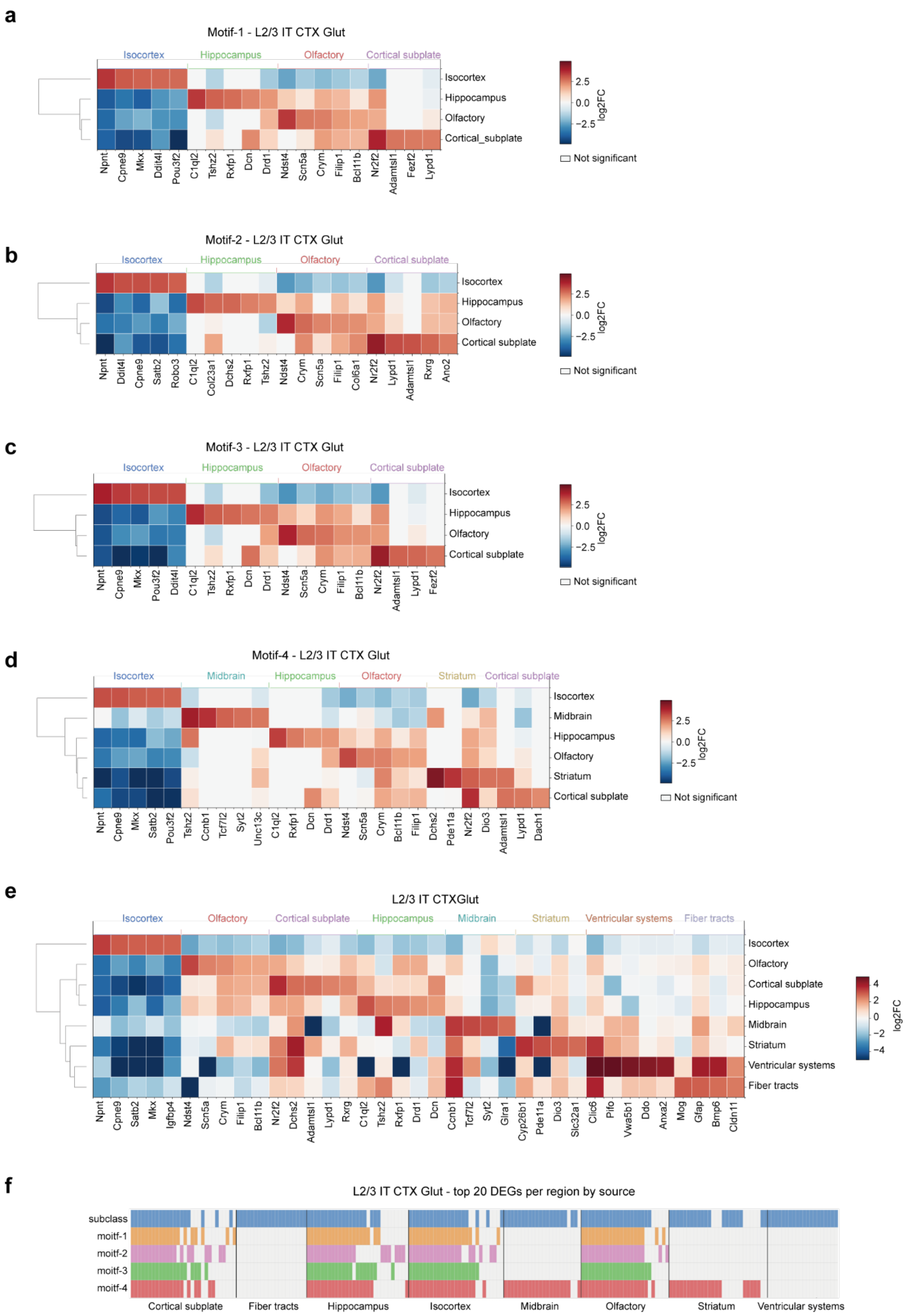


Supplementary Fig. 12 | Regional differential expression of motif+ glutamatergic neurons: L2/3 IT CTX Glut subclass. a–d, Top 5 differentially expressed genes per brain region for motif 1 (a), motif 2 (b), motif 3 (c), and motif 4 (d), controlling for subclass by restricting to L2/3 IT CTX Glut neurons. e, Subclass-level regional DEGs. f, Overlap between motif-context and subclass-level regional DEGs.

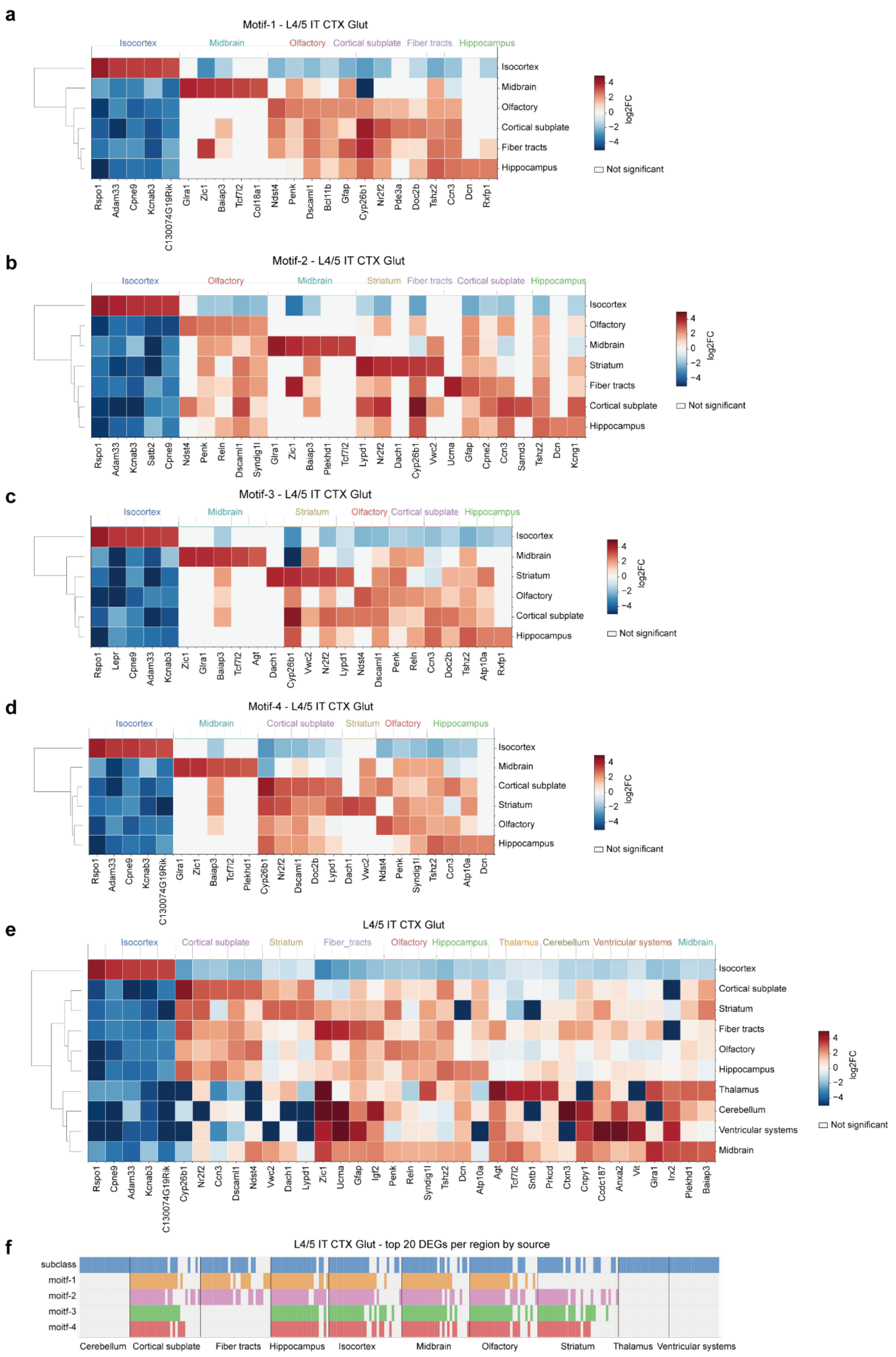


Supplementary Fig. 13 | Regional differential expression of motif+ glutamatergic neurons: L4/5 IT CTX Glut subclass. Same layout as Supplementary Fig. 15, for L4/5 IT CTX Glut neurons.

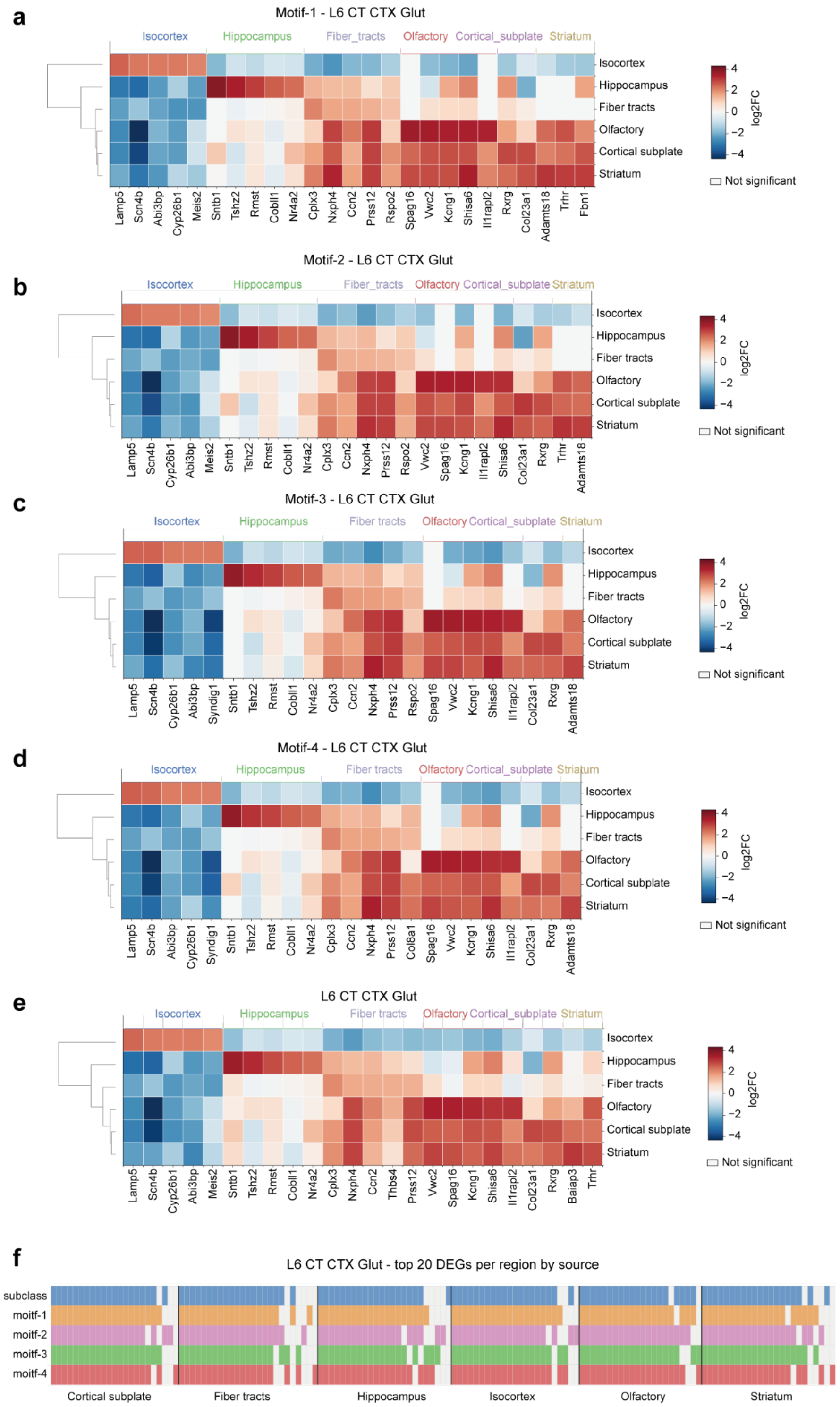


Supplementary Fig. 14 | Regional differential expression of motif+ glutamatergic neurons: L6 CT CTX Glut subclass. Same layout as Supplementary Fig. 15, for L6 CT CTX Glut neurons.

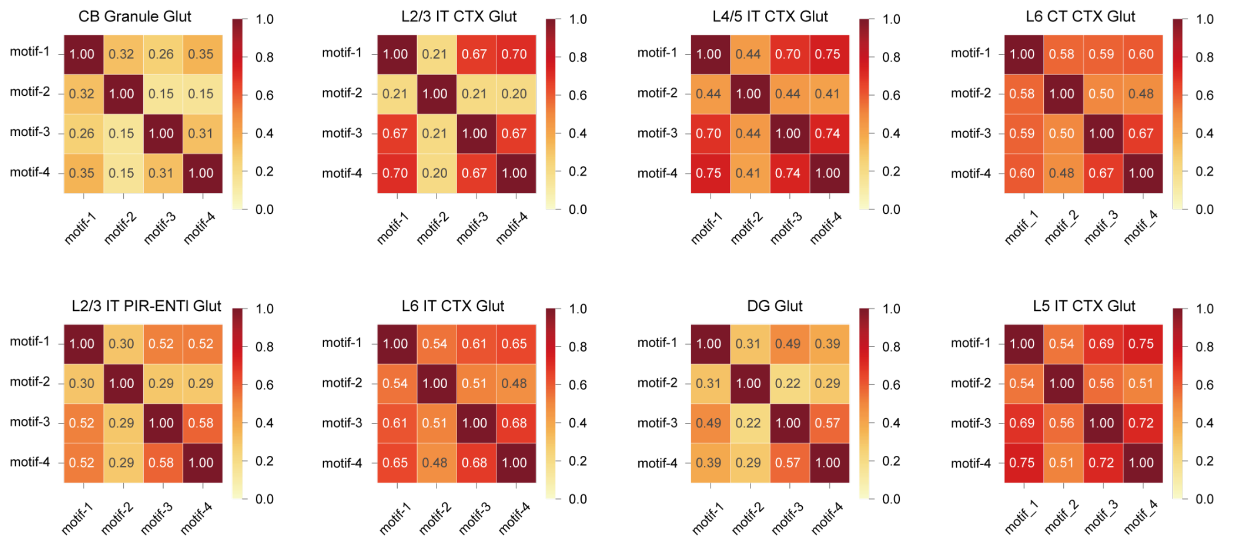


Supplementary Fig. 15 | Overlap of motif+ glutamatergic neurons across motifs within major subclasses. Pairwise Jaccard similarity of motif+ glutamatergic neuron populations across four motifs, computed separately for each of eight glutamatergic subclasses. Individual neurons frequently participate in multiple motif contexts concurrently.

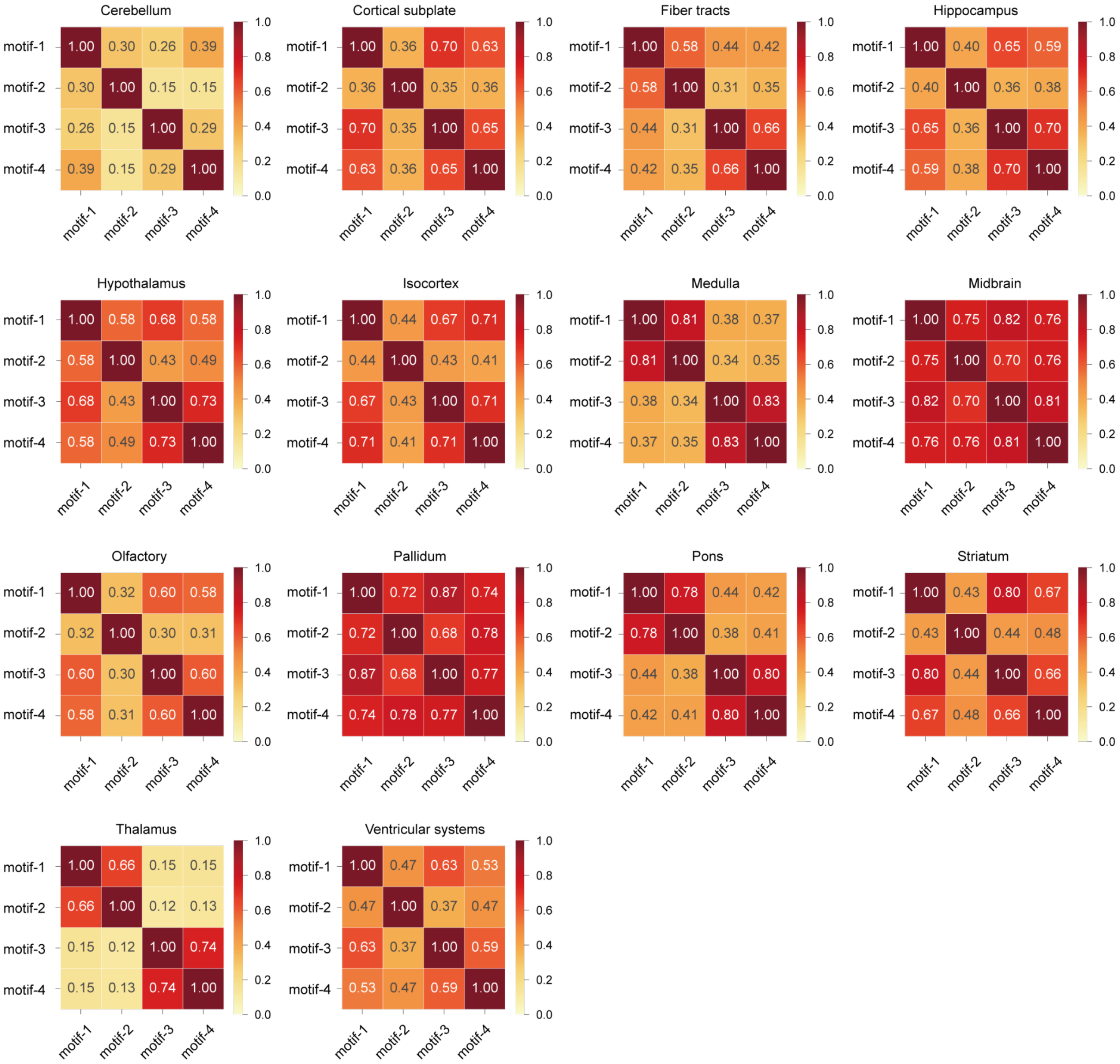


Supplementary Fig. 16 | Overlap of motif+ glutamatergic neurons across motifs within brain regions. Pairwise Jaccard similarity of motif+ glutamatergic neuron populations across four motifs, computed separately within each brain region.

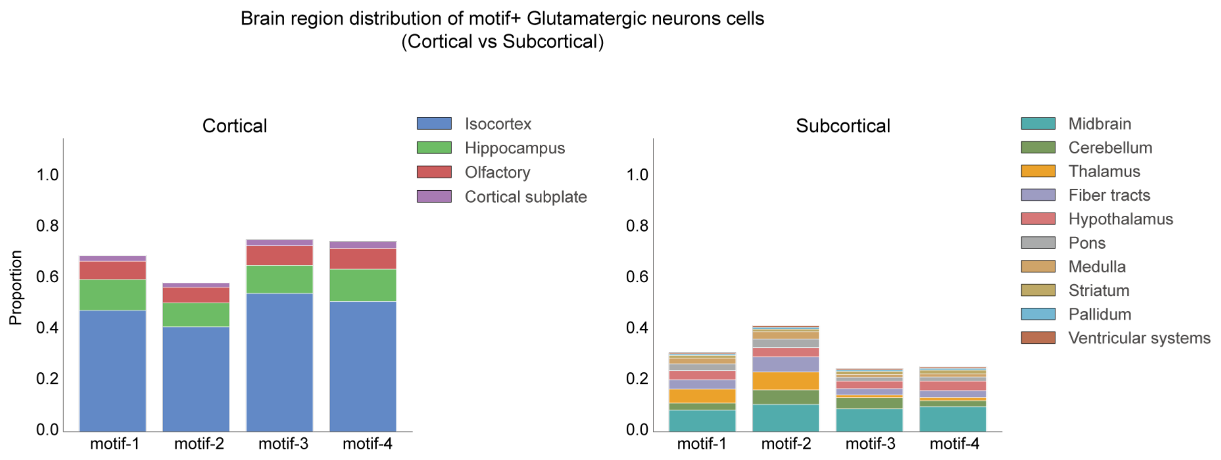


Supplementary Fig. 17 | Brain region distribution of motif+ glutamatergic neurons. Proportion of motif+ glutamatergic neurons residing in cortical (left) and subcortical (right) brain regions for each motif. Isocortex is the predominant region across all four motifs.

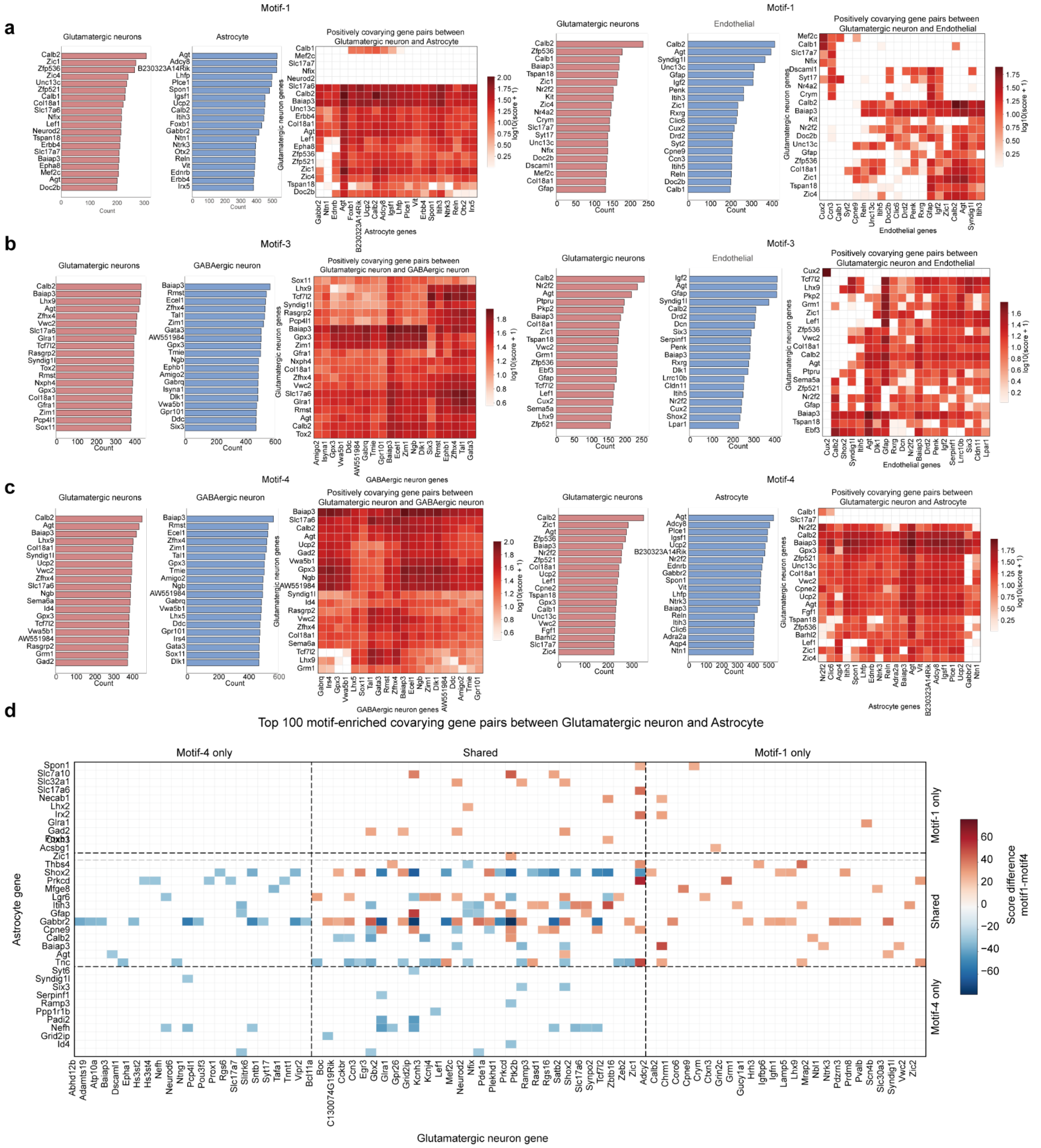


Supplementary Fig. 18 | Whole-brain covarying gene pairs for motifs 1, 3, and 4. a–c, For each motif, left panels show top 20 most frequently appearing anchor (glutamatergic neuron) and motif genes among positively covarying gene pairs; right panels show the corresponding covarying gene pair matrices. a, Motif 1 [glutamatergic, astrocyte, endothelial]. b, Motif 3 [glutamatergic, GABAergic, endothelial]. c, Motif 4 [glutamatergic, GABAergic, astrocyte]. d, Top 100 motif-enriched covarying gene pairs between glutamatergic neurons and astrocytes, comparing motif 1 versus motif 4. Gene pairs are grouped into motif-4-only, shared, and motif-1-only categories. Score difference (motif 1 minus motif 4) is shown.

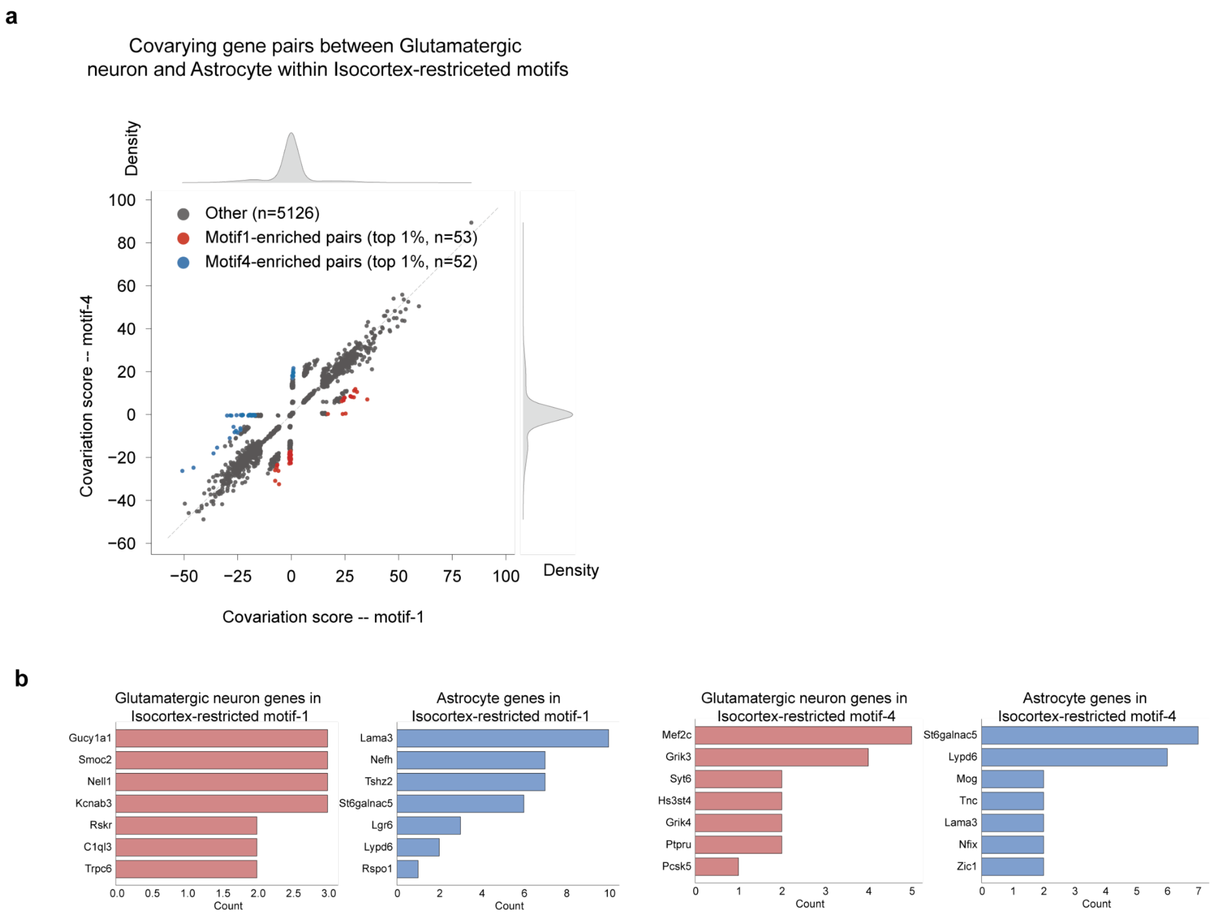


Supplementary Fig. 19 | Isocortex-restricted motif-associated covariation analysis. a, Scatter plot of covariation scores for glutamatergic neuron–astrocyte gene pairs in motif 1 (x-axis) versus motif 4 (y-axis) within isocortex. Top 1% motif-enriched pairs are highlighted for motif 1 (red) and motif 4 (blue). b, Top frequent anchor (glutamatergic neuron, red) and motif (astrocyte, blue) genes among isocortex-restricted motif-enriched covarying gene pairs, for motif 1 (left) and motif 4 (right).

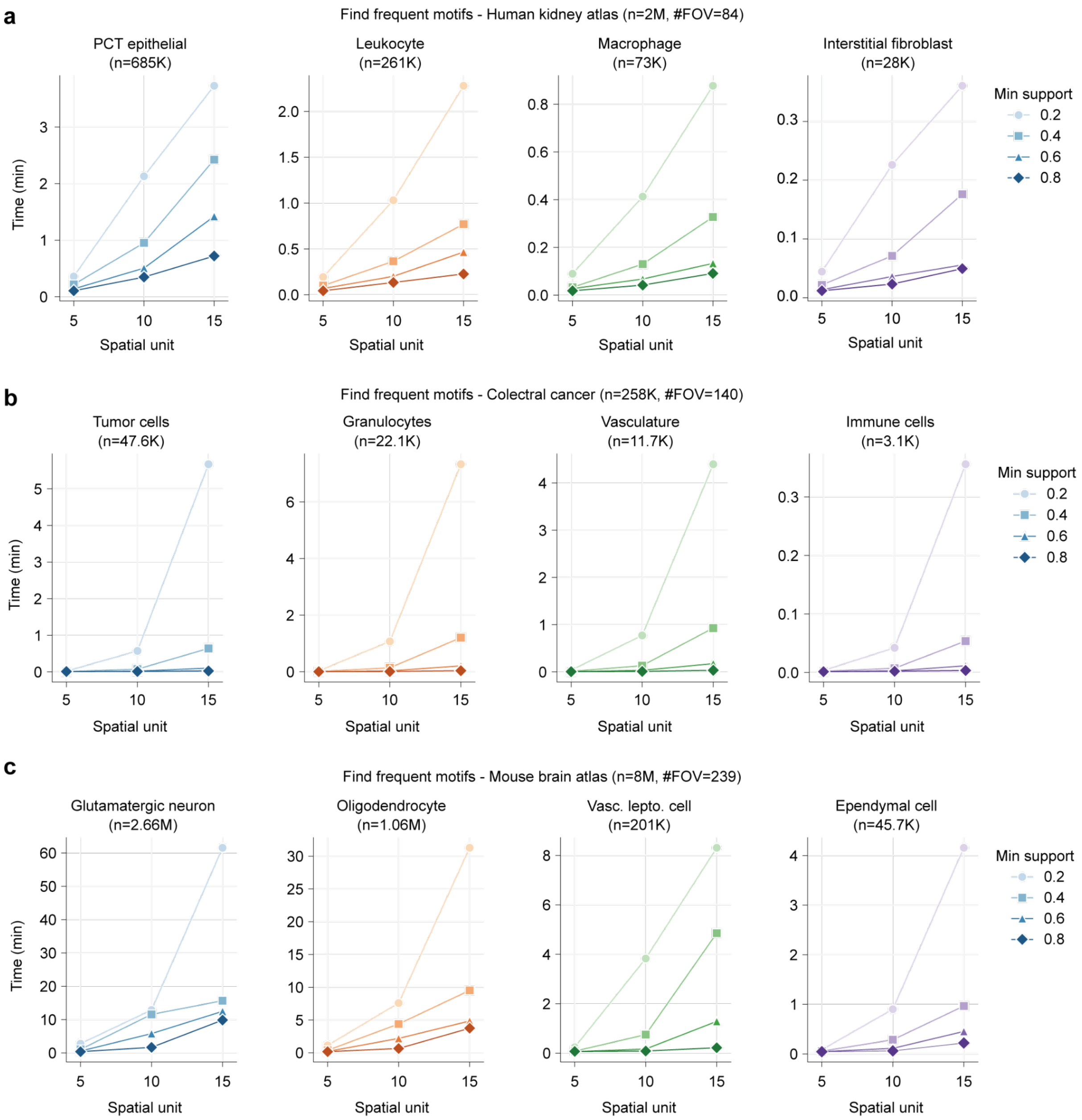


Supplementary Fig. 20 | Scalability of frequent motif identification. Runtime (minutes) for motif discovery (FP-Growth) as a function of neighborhood radius (5, 10, 15 spatial units) across three datasets: a, kidney atlas (~2M cells, 84 FOVs), b, colorectal cancer CODEX (~258K cells, 140 FOVs), and c, MERFISH mouse brain atlas (~8M cells, 239 FOVs). Within each dataset, four representative anchor cell types of varying abundance are shown. Lines indicate different minimum support thresholds (0.2, 0.4, 0.6, 0.8).

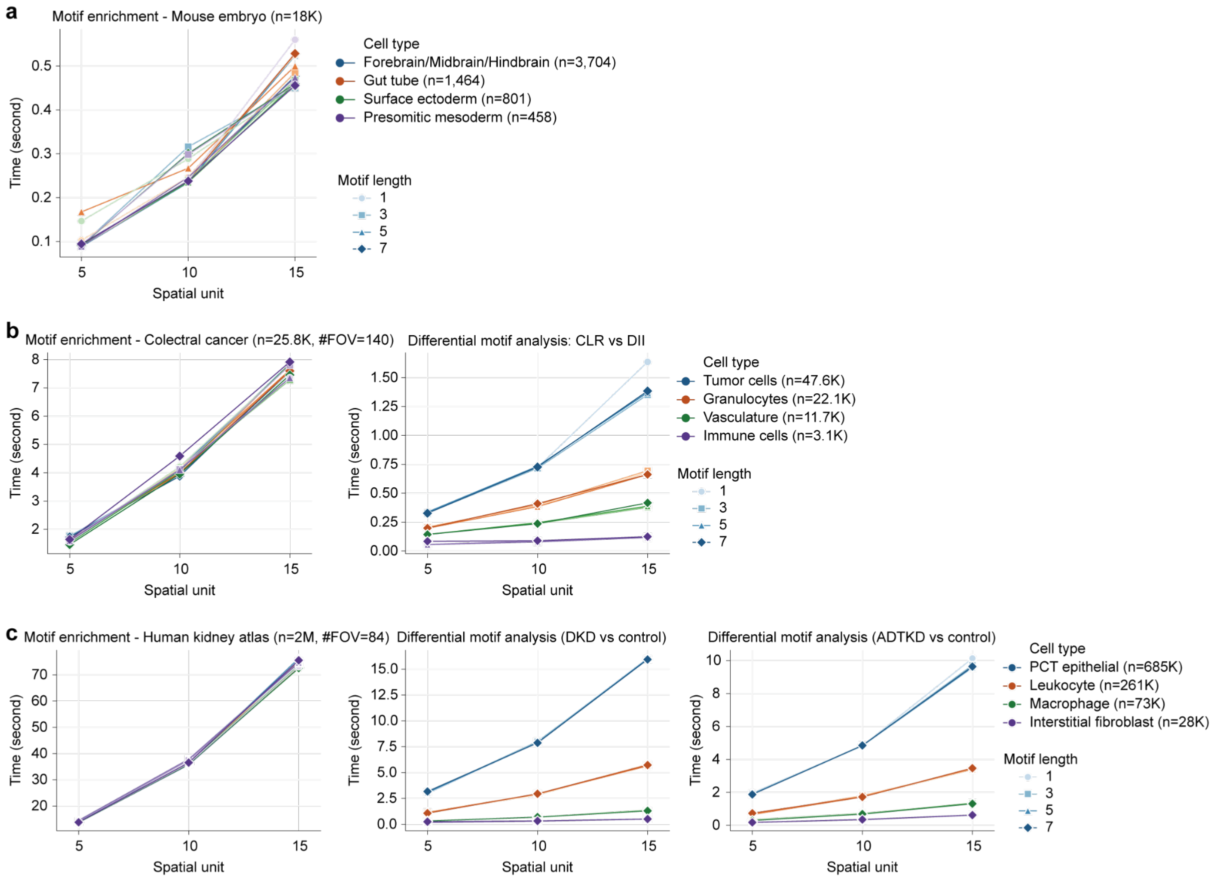


Supplementary Fig. 21 | Scalability of motif enrichment and differential motif analysis. Runtime (seconds) for motif enrichment analysis and differential motif analysis as a function of neighborhood radius and motif length. a, Motif enrichment in the mouse embryo dataset (~18K cells). b, Motif enrichment (left) and differential motif analysis between CLR and DII (right) in the colorectal cancer dataset (~258K cells). c, Motif enrichment (left) and differential motif analysis for DKD versus control (middle) and ADTKD versus control (right) in the kidney atlas (~2M cells).

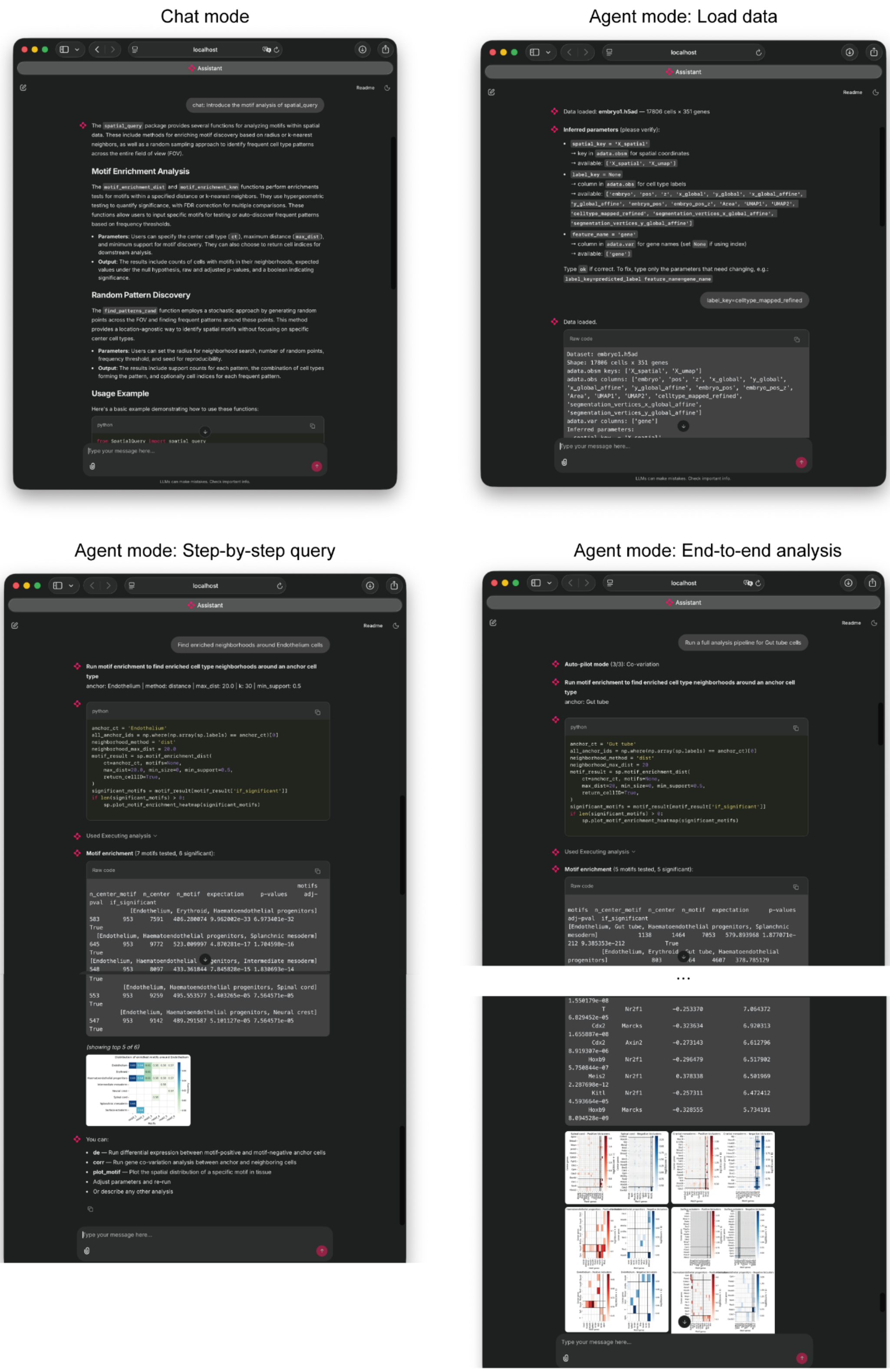


Supplementary Fig. 22 | Representative use cases of the SpatialQuery natural language agent. Screenshots of the Chainlit-based interface. Top left, chat mode returns method explanations and usage examples about SpatialQuery. Top right, agent mode with initialization of python objects: upon loading an .h5ad file, the agent infers key parameters from the AnnData object and prompts the user to confirm. Bottom left, step-by-step analysis in agent mode: the agent executes a user-specified analysis, returns results and visualization, and suggests downstream analyses. Bottom right, automated pipeline mode: a single instruction triggers sequential execution of motif enrichment, differential expression, and covariation analyses around specified anchor cell type.

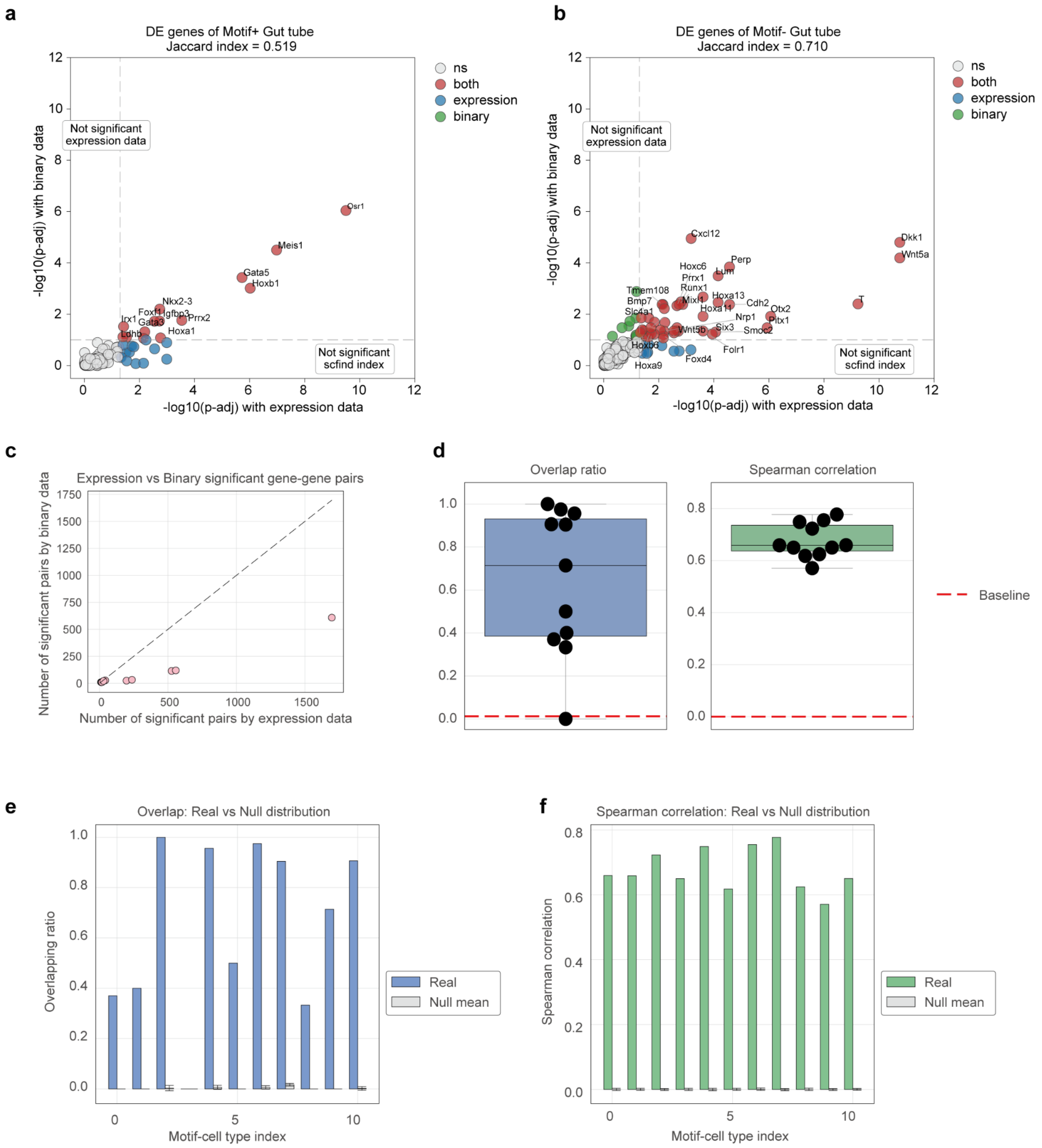


Supplementary Fig. 23 | Consistency between binary-based and expression-based analyses. a–b, Scatter plots comparing adjusted p-values of differentially expressed genes between binary (scfind index, significance threshold alpha = 0.1) and expression-based (alpha = 0.05) methods for motif+ (a) and motif− (b) gut tube cells. Jaccard indices indicate overlap of significant genes. c, Number of significant covarying gene pairs identified by each method across anchor–motif cell type combinations. The motif refers to enriched motifs surrounding gut tube (radius = 8, frequency threshold = 0.2), yielding 5 motifs and 11 cross-type anchor–motif cell type combinations. Each data point represents one combination. d, Overlap ratio (left) and Spearman rank correlation (right) of significant covarying gene pairs between binary and expression-based analyses, summarized across all 11 combinations. Each dot represents one anchor–motif cell type combination. Red dashed line indicates the mean of the permutation-derived null distribution. e–f, Per-combination comparison of overlap ratio (e) and Spearman correlation (f) between observed values and null distributions. For each combination, observed values (colored bars) are shown alongside the null mean (white bars).

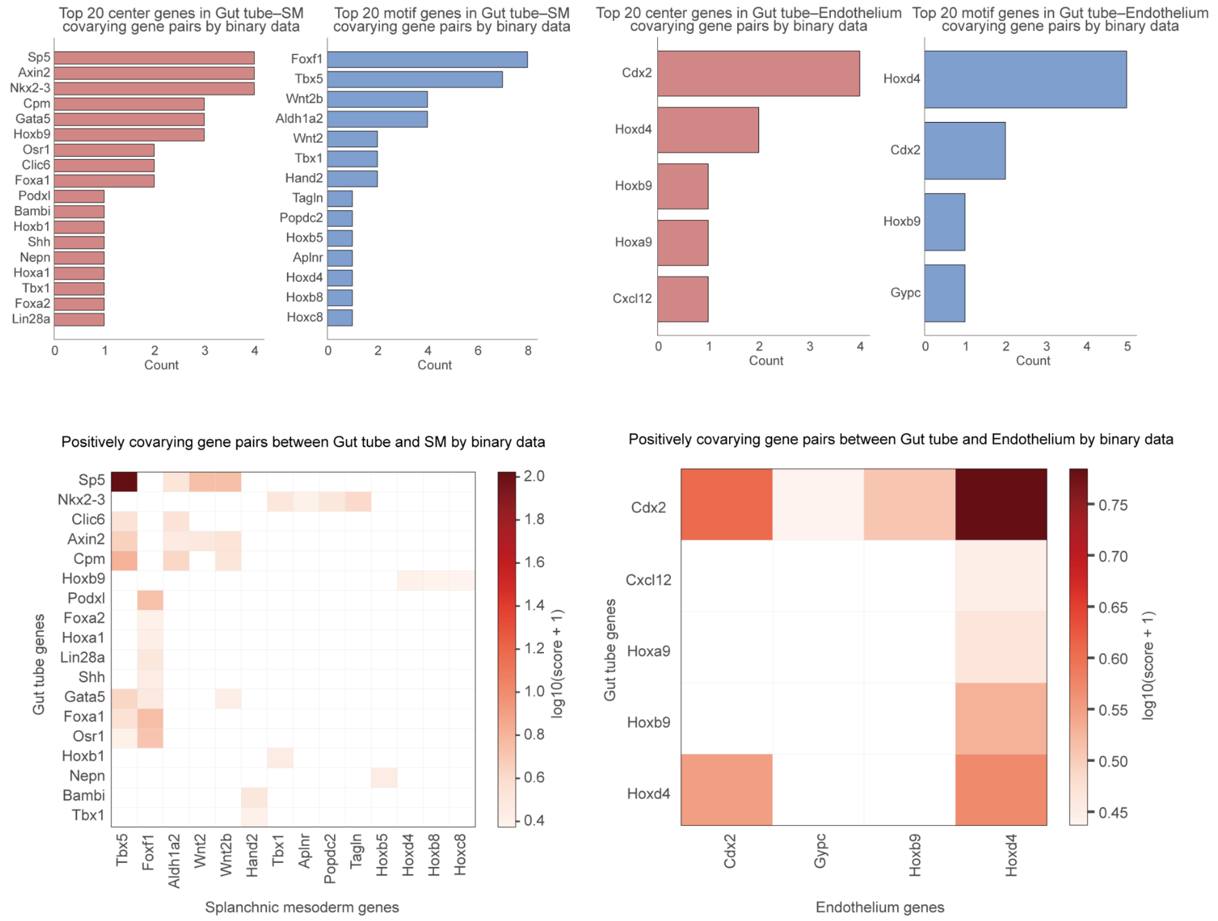


Supplementary Fig. 24 | Covarying gene pairs identified by binary-based analysis. Top, distribution of the top 20 anchor (gut tube) and motif genes among positively covarying gene pairs identified using scfind-indexed binary expression data, for gut tube–splanchnic mesoderm (left) and gut tube–endothelium (right) within [Splanchnic mesoderm–Endothelium] motif context. Bottom, corresponding covarying gene pair matrices involving the top genes.
